# Cross-species analyses reveal RORγt-expressing dendritic cells are a lineage of antigen presenting cells conserved across tissues

**DOI:** 10.1101/2024.05.06.592772

**Authors:** Hamsa Narasimhan, Maria L. Richter, Ramin Shakiba, Nikos E. Papaioannou, Christina Stehle, Kaushikk Ravi Rengarajan, Isabel Ulmert, Vanessa Küntzel, Eva-Lena Stange, Alina U. Antonova, Ludger Klein, Diana Dudziak, Marco Colonna, Natalia Torow, Mathias W. Hornef, Katharina Lahl, Chiara Romagnani, Maria Colomé-Tatché, Barbara U. Schraml

## Abstract

Conventional dendritic cells (cDCs) are potent antigen presenting cells (APCs) that exhibit tissue and age-specific diversity allowing them to direct situation-adapted immunity. Thereby they harbor great potential for being targeted in vaccination and cancer. Here, we resolve conflicting data about expression of retinoic acid receptor-related orphan receptor-γt (RORγt) in cDCs. We show that RORγt^+^ DCs exist in murine lymphoid and non-lymphoid tissues across age. Fate mapping, functional assays and single cell multiomic profiling reveal these cells as ontogenetically and transcriptionally distinct from other well characterized cDC subtypes, as well as from RORγt^+^ type 3 innate lymphocytes (ILC3s). We show that RORγt^+^ DCs can migrate to lymph nodes and activate naïve CD4^+^ T cells in response to inflammatory triggers. Comparative and cross-species transcriptomics revealed homologous populations in human spleen, lymph nodes and intestines. Further, integrated meta-analyses aligned RORγt^+^ DCs identified here with other emerging populations of RORγt^+^APCs, including R-DC-like cells, Janus cells/extrathymic Aire expressing cells (eTACs) and subtypes of Thetis cells. While RORγt^+^APCs have primarily been linked to T cell tolerance, our work establishes RORγt^+^ DCs as unique lineage of immune sentinel cells conserved across tissues and species that expands the functional repertoire of RORγt^+^ APCs beyond promoting tolerance.

**One sentence summary:** RORγt^+^ DC exhibit versatile APC functions and are a distinct immune lineage conserved across age, tissues and species that entails Thetis cells, Janus cells/RORγt^+^ eTACs and R-DC-like cells.

## INTRODUCTION

cDCs are immune sentinels located in lymphoid and non-lymphoid tissues (*1–3*). They effectively sense pathogens and subsequently migrate to and initiate T cell responses in secondary lymphoid organs (*1–3*). Their functional versatility makes them attractive for being targeted in vaccination against pathogens or cancer (*2–6*). Accordingly, considerable work has been invested in understanding the functional and ontogenetic diversity of cDCs, yet these cells remain ambiguous to define as fate mapping has recently uncovered novel populations with overlapping phenotype but distinct origin (*2*, *7–12*).

cDCs are generally considered to descend from committed myeloid cDC progenitors (*13–15*) and exist as functionally specialized subtypes. Type 1 cDCs (cDC1) excel at driving CD8^+^ T cell and T helper 1 (Th1) responses while cDC2 better promote Th2 and Th17 CD4^+^ T cell responses (*1–3*). cDC2 entail Notch2-dependent ESAM^high^ or T-bet-expressing cDC2A and Notch2-independent ESAM^low^ or T-bet-negative cDC2B (*16–18*). Both subtypes can induce Th17 responses but cDC2A appear more potent inducers of T follicular helper cell (Tfh) responses (*19*), while cDC2B may better promote Th2 responses (*20*). Fetal liver lymphoid progenitors in neonatal spleen and lymphoid derived transitional DCs (tDCs) in adult spleen can differentiate into cells resembling cDC2A and cDC2B (*9–11*, *21*). tDCs are cells with features of both cDCs and plasmacytoid DCs (pDCs), the latter of which are potent producers of type I interferons in response to viruses (*9*, *10*, *21*). Monocytic progenitors also generate cDC2B-like cells, which have been termed DC3 because of their distinct origin and similarity to human DC3 populations and in vitro appear superior in polarizing IL17A producing Th17 cells to cDC2B (*12*, *22*, *23*). The above heterogeneity therefore needs to be better defined to explore the full functional spectrum of cDCs.

In addition to cDCs, RORγt-expressing APCs have emerged as potent regulators of T cell tolerance (*24–34*). RORγt is a transcription factor encoded by the RORC gene that differs from its isoform RORγ by three amino acids at the amino terminus (*35*, *36*). RORγt-expressing APCs include subsets of ILC3s, RORγt^+^ extrathymic AIRE expressing cells (eTACs - including “Janus” and AIRE^+^ ILC3-like cells), as well as other surfacing populations like Thetis cells (*32–35*, *37*, *38*). Their lineage relationships and functional specializations are ill-defined because RORγt-expressing APCs phenotypically resemble each other and share expression of CD11c and ZBTB46 with cDCs (*29*, *33*, *34*, *38*, *39*). Janus cells express AIRE, RORγt and the integrin β 8 (ITGB8) but lack ILC3 markers CXCR6 and IL7R. They phenotypically and transcriptionally resemble CCR7^+^ migratory cDCs (*33*, *38*) and have primarily been profiled from pooled cervical, brachial, axillary, inguinal, and mesenteric lymph nodes (mLN) (*38*). Thus, it is unclear if they exhibit site-specific heterogeneity. Thetis cells have been described in neonatal mLN and strongly resemble Janus cells although they are apparently absent from skin draining LN (*34*). Despite their transcriptional similarities to Janus cells, only a fraction of Thetis cells (TCI and TC III) express AIRE, while TCIV expresses ITGβ8 but lacks AIRE (*34*) and it is unclear if Thetis cells even exist in adult mice (*40*). While Janus and Thetis cells in mice have been shown to drive T cell tolerance, R-DC-like cells that resemble Thetis cells have been described in human tonsil, where they can activate T cells (*41*). Thus, the above cell types may function beyond T cell tolerance, highlighting the need for a more accurate alignment of Janus cells, Thetis cells and RORγt^+^ eTAC subsets, and the tissue specific signals that regulate them.

cDC2B from adult mouse spleen show accessible chromatin at ROR response elements (RORE) in bulk ATAC sequencing, leading to the notion that cDC2B express RORγt (*16*, *35*). However, active RORγt protein expression in cDC2B from adult mice has not been reported and only a minor fraction of cDC2B show RORC expression history (*11*, *16*). In contrast to this, we have demonstrated active RORγt protein expression in a fraction of cDC2-like cells in neonatal mouse spleen and Peyer’s patches that phenotypically and transcriptionally more closely resemble cDC2A than cDC2B (*11*, *42*). These RORγt-expressing cDC2-like cells in neonatal spleen do not arise from *Clec9a*-expressing myeloid progenitors (*11*), raising the question whether RORγt-expressing cDC2-like cells are a unique type of APC, possibly related to Thetis cells, some subsets of which resemble cDC2 in terms of CD11b expression (*34*).

Here, we set out to reconcile conflicting reports regarding RORγt expression in cDCs to gain a better understanding of the functional heterogeneity, lineage relationships and phenotypic overlap of cDCs and RORγt^+^ APCs, which is a prerequisite to explore the specific immune functions of these cells. We show that RORγt^+^ DC that are ontogenetically and transcriptionally distinct from other cDC subtypes exist across tissues and species. These cells bear hallmark features of DCs, including developmental dependence on FLT3L, migration to lymph nodes and ability to activate naïve CD4^+^ T cells in response to inflammatory stimuli. Comparative transcriptomics further demonstrate that various previously described RORγt^+^ APC populations, including RORγt^+^ eTACs, Janus and Thetis cells, can be reconciled under the umbrella of RORγt^+^ DCs.

## RESULTS

### RORγt^+^ DC exist in murine spleen across age

To gain further insight into RORγt-expressing APCs in murine spleen we first profiled spleens from *RORγt^GFP^Clec9a^Cre^Rosa^Tom^*mice systematically across age by flow cytometry. In these mice Tomato expression tracks cells arising from committed *Clec9*a-expressing myeloid cDC progenitors (*14*), while GFP expression reports the RORγt-specific isoform encoded by RORC (*43*). We first identified MHCII^+^ILC3s as CD90^+^CD127^+^MHCII^+^GFP^+^ cells and then gated cDCs as CD11c^+^MHCII^+^ cells lacking the canonical ILC3 markers CD90 and CD127 - also known as interleukin-7 Receptor (IL-7R) (Fig. 1a, Fig. S1a). Within CD11c^+^MHCII^+^ splenocytes, we detected GFP expressing cells at all ages examined (Fig. 1a,b, Fig. S1a). Importantly, CD11c^+^MHCII^+^ cells which were GFP^+^ also stained with an anti-RORC antibody (Fig. S1b). These data confirm that GFP signal in *RORγt^GFP^*mice accurately reports RORγt protein expression and indicates that anti-RORC antibody staining in CD11c^+^MHCII^+^ cells predominantly identifies cells expressing the γt isoform of RORC. Thus, RORγt-expressing cDC like cells exist in murine spleen across age and these cells can either be detected using an anti-RORC antibody or by GFP signal in *RORγt^GFP^* mice (Fig. 1a, S1b). Unbiased gating of RORγt^+^ cells showed that RORγt^+^CD11c^+^MHCII^+^ cells lacked ILC3 markers CD127, CD90 and CXCR6, establishing them as phenotypically distinct from MHCII^+^ ILC3 (Fig. S1c, e, f). The frequency of RORγt^+^ cells within CD11c^+^MHCII^+^ cells declined with age, however, their absolute numbers increased until weaning age, correlating with organ growth and from weaning numbers remained relatively constant until adulthood (8-15 weeks of age) (Fig. 1d, Fig. S1d).

**Figure 1.**
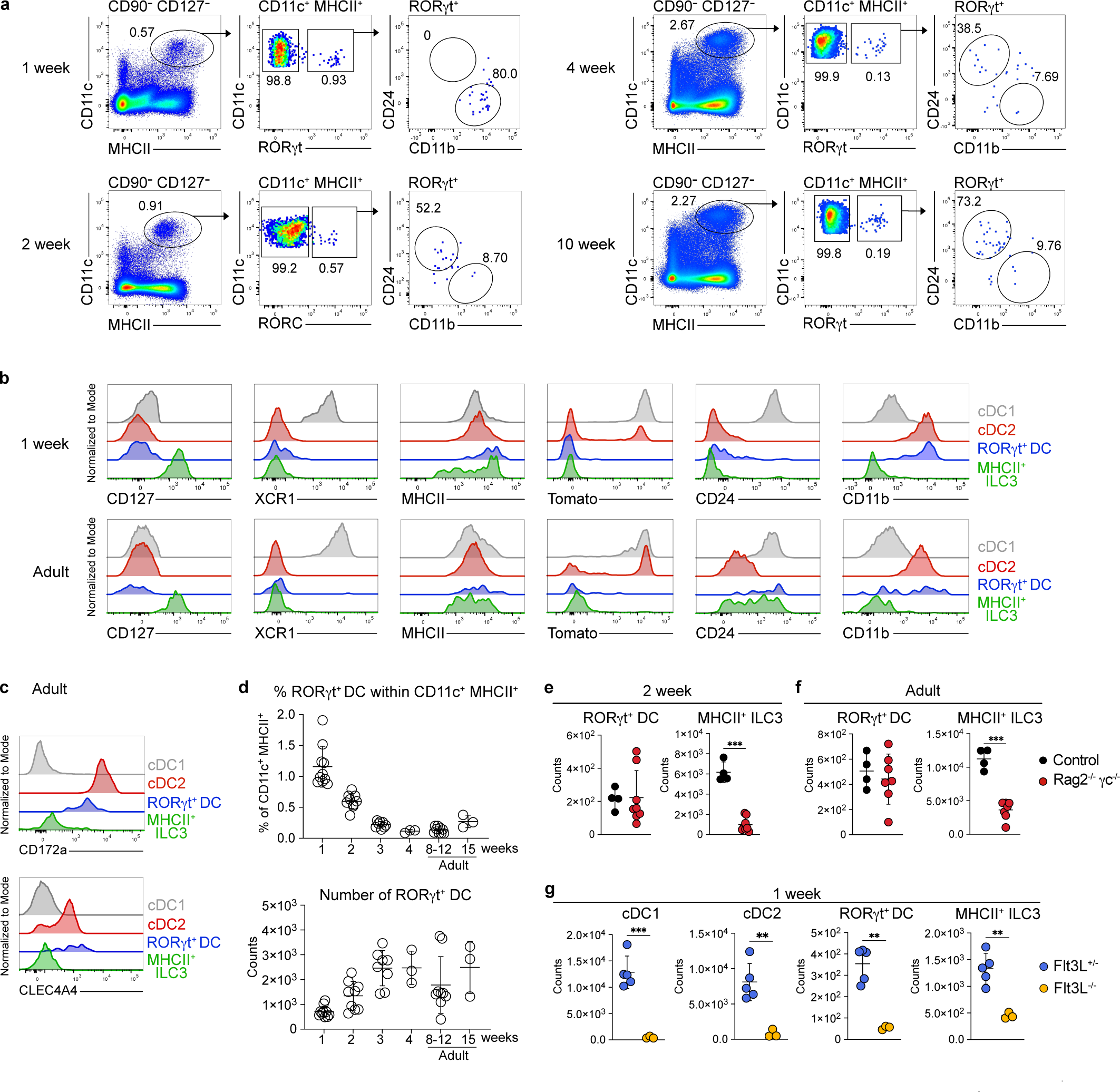
RORγt^+^ DCs exist in murine spleen across age. **a, b.** Splenocytes from *Clec9a^Cre^Rosa^Tom^* or *Clec9a^Cre^Rosa^Tom^RORγt^GFP^*mice at the indicated ages were analyzed by flow cytometry **a.** Live CD90^-^CD127^-^CD11c^+^MHCII^+^ cells were gated and RORγt^+^ cells revealed by GFP expression or anti-RORC intranuclear staining (2-week time point). GFP^+^ or RORC^+^ cells were further analyzed for CD11b and CD24 expression. **b,c.** Expression of the indicated surface markers in cDC1 (grey), cDC2 (red), MHCII^+^ ILC3 (green) and CD11c^+^MHCII^+^RORγt^+^ cells (blue) from spleens of 1-week-old and adult mice. Gating strategy see Figure S1a. **d**. Frequency of RORγt^+^ cells within CD11c^+^MHCII^+^ cells and number of RORγt^+^ DCs in spleens of the indicated ages (1-week-old n=10; 2-week-old, n=9; 3-week-old n=8, 4-week-old n=3; 8-12-week-old n=6; 15-week-old n=3). Data are pooled from 1-2 independent experiments; each data point represents a biological replicate. **e, f.** MHCII^+^ ILC3 and RORγt^+^ DCs were quantified in spleens from 2-week-old **(e)** and adult *Rag2*^-/-^*γc*^-/-^ mice **(f)** and littermate controls. Each data point represents an individual mouse from two independent experiments. **g.** Quantification of cDC1, cDC2, RORγt^+^ DC, and MHCII^+^ ILC3 in spleens from 1-week-old *Flt3*^-/-^ and *Flt3*^+/-^ littermate controls. RORγt^+^ DC quantified by intranuclear staining against RORC. Each dot represents one mouse, horizontal bars represent mean, error bars represent SD. **p (0.0021) ***p (0.0002). Statistical analyses in (**e-g**) were performed using two-tailed Welch’s *t*-test.

This observation was surprising, since we had previously found that RORγt^+^ cDC2-like cells were nearly undetectable by two weeks of age (*11*). Thus, we examined the phenotype of CD11c^+^MHCII^+^RORγt^+^ cells across age. At one week of age CD11c^+^MHCII^+^RORγt^+^ cells uniformly expressed CD11b but lacked CD24, however, starting at two weeks of age RORγt^+^ cells showed a CD11b^low^ to negative phenotype and stained positive for CD24 (Fig. 1a and b). As observed in neonates (*11*), RORγt^+^CD11c^+^MHCII^+^ cells from adult spleen expressed the cDC2 markers CD172a and CLEC4A4 (Fig. 1c). At all ages examined RORγt^+^CD11c^+^MHCII^+^ cells stained negative for the cDC1-specific marker XCR1 (Fig. 1b). Thus, the apparent lack of CD11b by two weeks of age explains why these cells had previously been missed in our analyses of cDC2 in adult mice and suggests that these cells are either heterogenous or change their phenotype with age. In line with our previous work, RORγt^+^CD11c^+^MHCII^+^ cells lacked Tomato expression in *Clec9a^Cre^Rosa^TOM^* mice at all ages examined, supporting that they do not arise from *Clec9a*-expressing progenitors (*11*, *14*). To address relatedness of RORγt^+^CD11c^+^MHCII^+^ cells to ILC3s, we profiled *Rag2^-/-^γc^-/-^* mice, which lack ILC3s and their precursors (*44*). In these mice MHCII^+^ ILC3s were strongly reduced as expected, while RORγt^+^CD11c^+^MHCII^+^ cells were unaltered compared to control mice (Fig. 1e-f). Mice deficient for fms-like tyrosine kinase 3 ligand (FLT3L), a growth factor critical for the development of all DCs and ILCs (*45*, *46*), showed a strong reduction of cDC1, cDC2, ILC3s and RORγt^+^CD11c^+^MHCII^+^ cells (Fig. 1g). These data suggest that RORγt^+^CD11c^+^MHCII^+^ cells, like cDCs and ILC3s, require FLT3L for their development but that they arise from distinct yet to be specified progenitors. Because RORγt^+^CD11c^+^MHCII^+^ cells phenotypically and transcriptionally (*11*) resemble cDCs we refer to these cells as RORγt^+^ DCs from hereon for simplicity in accordance with suggestions for cDC and ILC nomenclature (*1*, *35*, *47*).

### RORγt^+^ DCs are bona fide antigen presenting cells

RORγt^+^ DCs expressed similar levels of MHCII compared to cDC1 and cDC2 at all ages examined (Fig. 1b), suggesting that they are APCs. To address if RORγt^+^ DCs have the capacity to activate naïve T cells, which is a hallmark feature of cDCs (*2*), we sort-purified RORγt^+^ DCs from spleens of two-week-old and adult *RORγt^GFP^* mice pulsed them with Ovalbumin (OVA) peptide 323–339 (OVA_323–339_) and subsequently cultured them with cell trace violet (CTV)-labelled naïve OTII transgenic T cells from adult mice in the absence or presence of T-cell polarizing cytokines. cDC2 and MHCII^+^ILC3 from the same mice were used as controls (Fig. S2A). Because RORγt^+^ DCs are limited in number (Fig. 1d), 250 APCs were cultured with 2500 OTII T cells per condition. As expected, cDC2 from two-week-old and adult mice induced T cell proliferation under all conditions tested (Fig. 2a,e, Fig. S2b,c), while MHCII^+^ILC3s were poor inducers of OTII T cell proliferation (Fig. 2a,e S2b-g). Importantly, under all conditions tested RORγt^+^ DCs from two-week old and adult mice stimulated the proliferation of naïve T cells, although to a lower extent than cDC2, and supported the differentiation of OTII cells into effector T cells, as measured by intracellular cytokine and FOXP3 staining (Fig. 2b-h). While RORγt^+^ DCs supported the differentiation of FOXP3^+^ Tregs equally or better than cDC2, the induction of Th17 cells by RORγt^+^ DCs was reduced compared to cDC2 (Fig. 2c,d,g,h). Thus, RORγt^+^ DCs can activate naïve T cells and induce their effector differentiation albeit with qualitative and quantitative differences compared to cDC2. In contrast, MHCII^+^ ILC3 induced little if any T cell proliferation and may require stimulation with cytokines, such as IL1β, to achieve their full APC potential (*48*).

**Figure 2.**
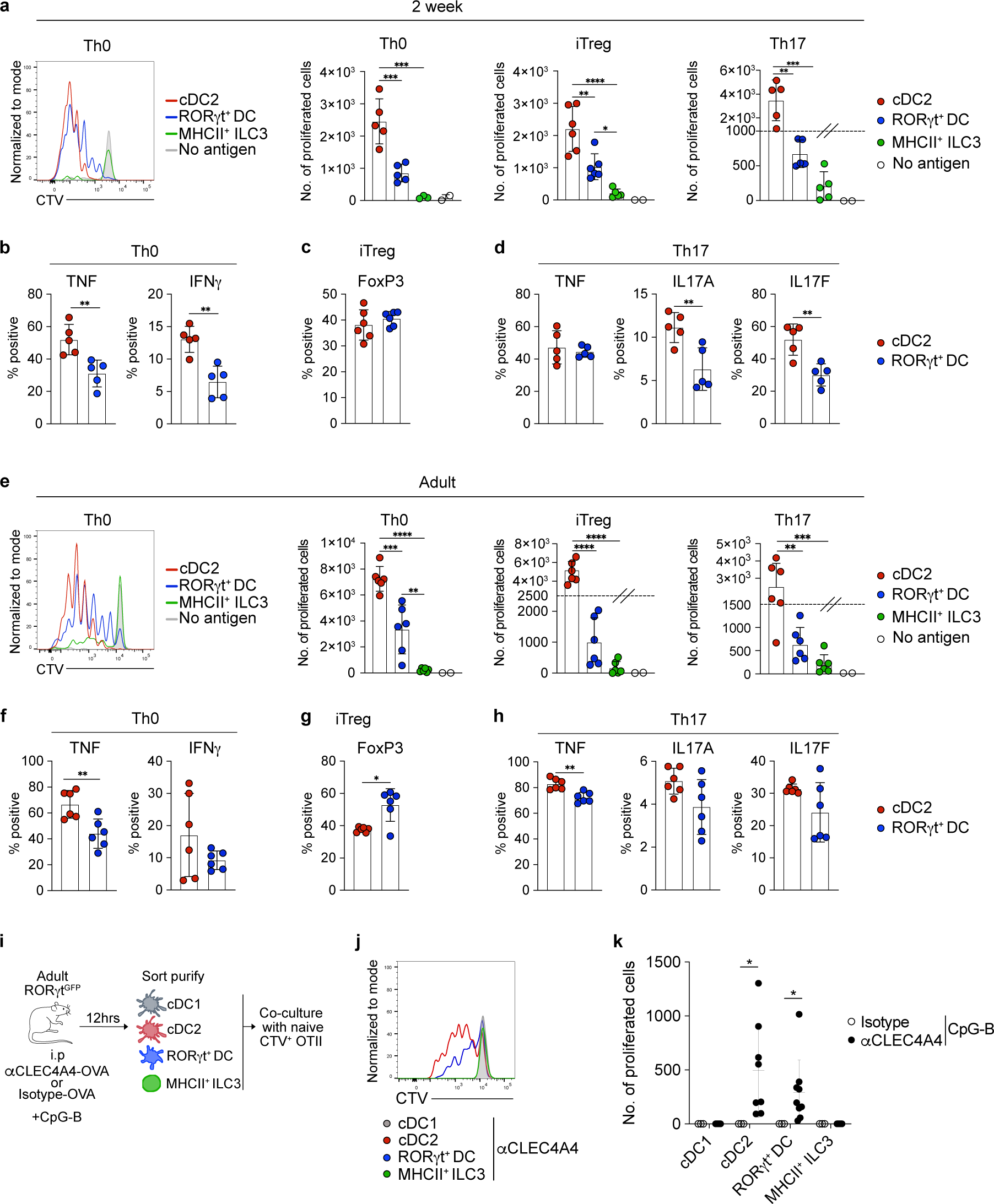
RORγt^+^ DCs are bona fide antigen presenting cells. **a-h.** cDC2, MHCII^+^ ILC3 and RORγt^+^ DCs from spleens of 2-week-old **(a-d)** or adult **(e-f)** *RORγt^GFP^* mice were pulsed with OVA_323-339_. 250 APCs were then co-cultured with 2500 CTV-labelled naïve OTII cells as indicated and 3.5 days later proliferation (CTV dilution), cytokine production and FOXP3 expression in proliferated T cells were quantified. **a.** CTV-trace of OTII cells co-cultured with cDC2 (red), RORγt^+^ DCs (blue), MHCII^+^ ILC3 (green), or cDC2 without OVA_323-339_ (grey) under non-polarizing conditions (Th0). Right: Quantification of total proliferated cells at end of culture with cDC2 (red), RORγt^+^ DCs (blue), MHCII^+^ ILC3 (green) or cDC2 without OVA_323-339_ (open circle) under the indicated conditions. **b-d.** Proliferated OTII cells were analyzed for TNF and IFNγ production **(b)**, FOXP3 expression **(c)**, or TNF, IL-17A and IL-17F production **(d)**. **e-h.** OTII T cells co-cultured with cDC2, MHCII^+^ ILC3 and RORγt^+^ DCs from adult mice were analyzed as in **a-f** above. **i-k.** Adult *RORγt^GFP^* mice were injected i.p. with anti-CLEC4A4-OVA or isotype-OVA control antibody plus CpG-B. After 12 hours 300 cDC2, cDC1, MHCII^+^ILC3 and RORγt^+^ DCs were sorted and co-cultured with 3000 naïve CTV-labelled OTII cells. **i.** Experimental set up. **j.** CTV dilution and **k.** number of proliferated OTII cells after co-culture with the indicated populations are shown (nβ5). Each dot represents one biological replicate from 2 independent experiments for each timepoint, horizontal bars represent mean, error bars represent SD. *p (0.0332), **p (0.0021) ***p (0.0002), ****p < 0.0001. Statistical analysis was performed using two-tailed Welch’s t-test **(b-d, f-h)** or one-way ANOVA with Tukey’s multiple comparisons **(a,e)**. Only statistically significant comparisons are indicated.

Having established that RORγt^+^ DCs have the potential to activate naïve T cells in vitro, we next asked if they also have the capacity to process and present antigen upon direct delivery of antigens in vivo. Coupling antigens to an antibody directed against the C-type lectin receptor CLEC4A4/DCIR2 allows to deliver antigens to cDC2 in vivo (*49*). Specific expression of CLEC4A4 on cDC2 and RORγt^+^ DCs from neonatal and adult mice (*11*), but not on cDC1 or MHCII^+^ ILC3 (Fig. 1c) (*49*), suggested that anti-CLEC4A4-OVA may allow to deliver antigen to RORγt^+^ DCs in vivo. We thus injected adult *RORγt^GFP^* mice with anti-CLEC4A4-OVA or isotype-matched control antibody in the presence of CpG-B as adjuvant, which strongly boosts T cell activation by cDC2 in this assay (*11*). 12 hours later, we sorted 300 cDC1, cDC2, RORγt^+^ DCs and MHCII^+^ILC3s and co-cultured them with 3000 naïve CTV-labelled OTII T cells (Fig. 2i-k). cDC2 and RORγt^+^ DCs induced OTII proliferation after targeting with anti-CLEC4A4-OVA but not isotype-matched control antibody (Fig. 2j,k), demonstrating successful targeting. In contrast, cDC1 and MHCII^+^ ILC3s did not induce T cell proliferation demonstrating cell type specific antigen targeting (Fig. 2j, k). OTII proliferation induced by RORγt^+^ DCs appeared marginally lower than that achieved by cDC2, which could be due to different efficiency of CLEC4A4 targeting or different responsiveness of RORγt^+^ DCs to CpG-B compared to cDC2. Taken together, these data demonstrate that RORγt^+^ DCs can process and present antigen and activate naïve CD4^+^ T cells in response to an inflammatory trigger, establishing RORγt^+^ DCs as bona fide APCs.

### Single cell multiomic profiling of the transcriptome and epigenome establishes the unique identity of RORγt^+^ DCs

To gain insight into the transcriptional and regulatory relatedness of RORγt^+^ DCs to cDCs and ILC3s, we performed paired single-cell RNA sequencing (scRNA-seq) and single-cell assay for transposase-accessible chromatin (scATAC) from the same cell using 10X multiomic profiling. To further assess if the transcriptional profile of RORγt^+^ DCs changes across age, RORγt^+^ DCs and MHCII^+^ ILC3s were sorted from spleens of two-week-old or adult mice and mixed at a 1:1 ratio (Fig. 3a) to enrich for RORγt^+^ DCs. CD11c^+^MHCII^+^ cDCs were added in 10-fold excess to capture their heterogeneity (Fig. 3a). After quality filtering, we retained chromatin accessibility and gene expression profiles from 9,899 and 11,980 nuclei respectively. Unsupervised clustering based on gene expression (RNA) and open chromatin (ATAC) profiles revealed 18 and 16 clusters, respectively (Fig. S3a, b). Based on published gene signatures (Table 1), these could be classified as cDC1, cDC2, migratory cDCs, ILC3s, pDCs and tDCs (Fig. 3b, Fig. S3a-f, Table 1). cDC2 split into 5 clusters based on gene expression, of which clusters 1, 2 and 3 scored high for cDC2A and ESAM^high^ signature genes (*16*, *50*), thus corresponding to cDC2A (Fig. S3c). Of note, cDC2 clusters 1 and 2 showed uniform chromatin accessibility (Fig. S3a, d), as discussed below. cDC2 clusters 4 and 5 transcriptionally resembled both cDC2B and DC3 (Fig. S3a,c,e) and could not confidently be delineated as either cell type based on published cell type specific markers (Fig. 3b, Fig. S3c, e) (*12*, *16*). We therefore refer to cDC2 clusters 4 and 5 as cDC2B/DC3. pDCs showed uniform chromatin accessibility but split into two clusters based on gene expression (Fig. S3a, b, d). *Rorc^+^* cells distributed across 5 clusters. Of these, clusters annotated as ILC3_1 and ILC3_2 scored high for ILC3 signature genes and expressed *Rora, Il7r, Cxcr6*. One cluster located close to ILC3_1 transcriptionally resembled ex-ILC3 (*51*) (Fig. 3b,c, Table 2). Additionally, one cluster expressed *Rorc*, *Prdm16,* and *Lingo4* (*11*) while lacking *Il7r, Rora*, and *Cxcr6* and showed significant similarity to RORγt^+^ DC2-like cells from neonatal mouse spleen (*11*), establishing it as RORγt^+^ DC (Fig. 3c,d, Table 1). We detected some *Aire* expression in migratory DCs (Fig. 3c), consistent with previous reports (*38*, *52*), but identified a distinct cluster of cells expressing *Aire, Rorc* and *Prdm16*, which we termed RORγt^+^ eTACs. RORγt^+^ DCs and RORγt^+^ eTACs transcriptionally most closely resembled each other and migratory cDCs (Fig. 3g)(*33*, *38*).

**Figure 3.**
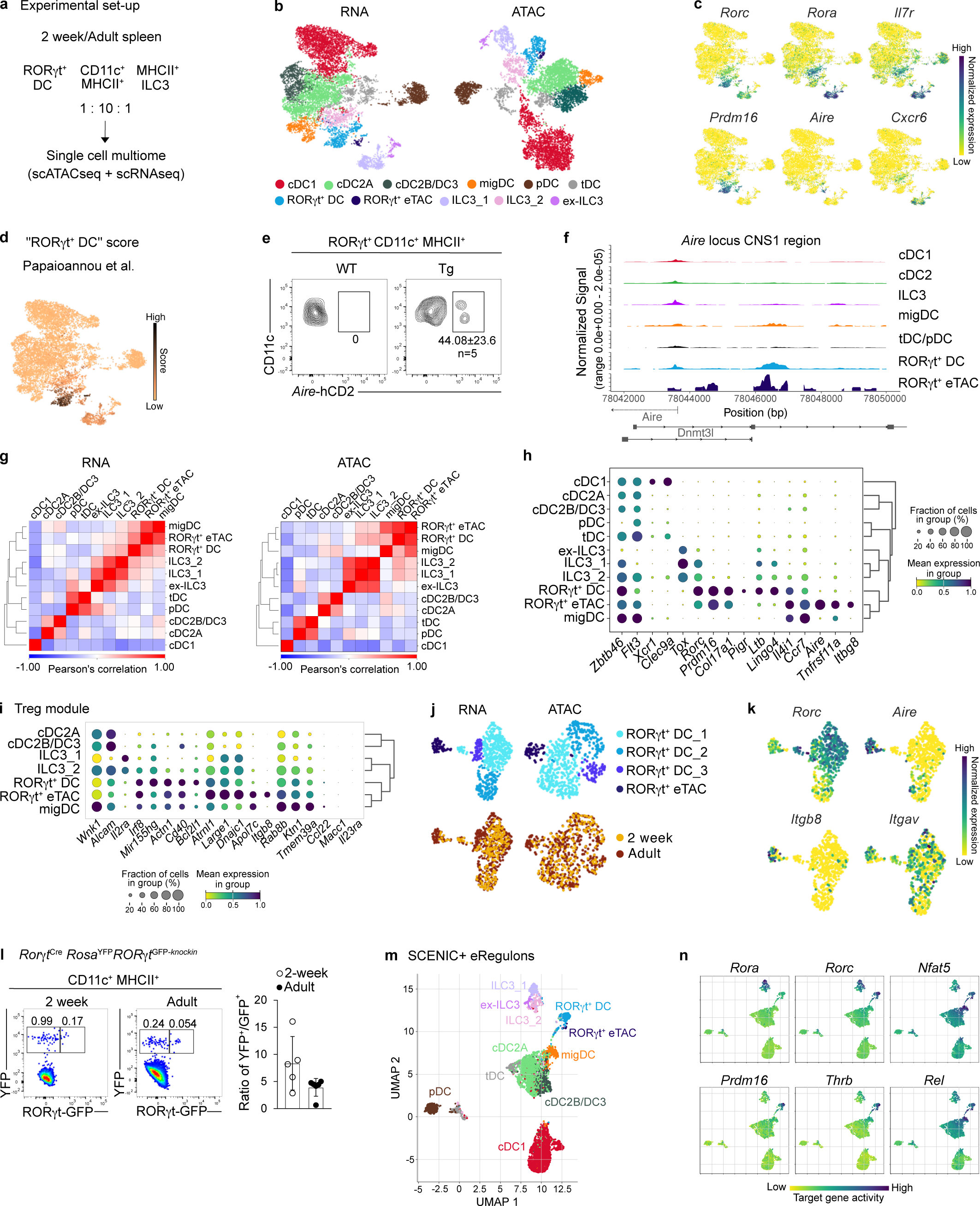
Single cell multiomic profiling reveals the unique identity of RORγt^+^ DCs. **a.** Experimental set up for single cell multiomic profiling (paired scRNA-seq and scATACseq) of RORγt^+^ DCs, RORγt^-^CD11c^+^MHCII^+^ cells and MHCII^+^ ILC3 from spleens of 2-week-old (n=2) or adult (n=3) *RORγt^GFP^* mice. **b.** RNA-based and ATAC-based UMAP of 11,980 nuclei annotated by cell type (see also Fig. S3). **c.** Expression of indicated genes on the RNA-based UMAP. **d.** RNA-based UMAP depicting enrichment score for genes that distinguished RORγt^+^ DC2-like cells in neonatal spleen (*11*). **e.** AIRE expression in RORγt^+^CD11c^+^MHCII^+^ cells revealed by anti-hCD2 staining in spleen of adult *Aire^hCD2^* mice. **f.** Chromatin accessibility at the CNS1 region of *Aire* locus in the indicated cell types. **g.** Pearson’s correlation to establish the similarity of individual clusters identified based on RNA and ATAC profiles. **h, i.** Bubble blot of select cell type defining genes **(h)** and genes involved in Treg induction **(i)**. **j.** RORγt^+^ DCs and RORγt^+^ eTACs were reclustered. The resulting UMAPs based on RNA and ATAC profiles are shown and annotated by timepoint below. **k.** Expression of *Rorc, Aire, Itgav* and *Itgb8* on the RNA-based UMAP from **(j)**. **l.** Spleens from 2-week-old and adult *RORγt*^Cre^*Rosa*^YFP^*RORγt*^GFP/wt^ mice were analyzed by flow cytometry. YFP (RORγt expression history) versus GFP (active RORγt) expression in CD11c^+^MHCII^+^ cells were plotted and the ratio of YFP^+^ to GFP^+^ cells (active RORγt expression) was calculated. Each dot represents one mouse. **m.** UMAP display of dimensionality reduction based on target genes and region enrichment scores generated using SCENIC+. **n.** UMAP colored by target gene activity of the indicated eRegulons predicted to regulate specific cell types (Fig. S4a).

Overall, the clusters identified based on chromatin accessibility and gene expression profiles were largely congruent (Fig. S3d). It was notable that RORγt^+^ eTACs and RORγt^+^ DC showed similar chromatin accessibility but somewhat diverged based on gene expression profile (Fig. 3b), raising the possibility that RORγt^+^ eTACs are a state of RORγt^+^ DC. Flow cytometry in *Aire^hCD2^* reporter mice (*53*) confirmed that a fraction of RORγt^+^ DCs indeed express AIRE (Fig. 3e). Accessibility mapping of the cis-regulatory region of the Aire locus, including the conserved noncoding sequence 1 (CNS1) region (*38*), confirmed accessible chromatin peaks in migDCs, RORγt^+^ DCs and RORγt^+^ eTACs. Highest accessibility of this region in the *Aire* locus was observed in RORγt^+^ DCs and RORγt^+^ eTACs and some cell type specific peaks were observed close to the transcriptional start site of *Aire* in RORγt^+^ eTACs (Fig. 3f). Differential gene expression analyses showed that the RORγt^+^ DC cluster was distinguished from other clusters by expression of genes including *Prdm16, Col17a1, Pigr, Ltb* and *Lingo4*, of which *Prdm16* and *Col17a1* expression were shared with RORγt^+^ eTACs (Fig. 3h, Table 2). RORγt^+^ eTACs showed specific expression of *Aire, Tnfrsf11a* (RANK) and *Itgb8*, establishing them as a likely equivalent of Janus cells/eTACs found in lymph nodes (Fig. 3h) (*33*, *37*, *38*). Importantly, the cDC associated marker *Zbtb46* was expressed on RORγt^+^ DC, RORγt^+^ eTACs, ILC3, tDC and cDC clusters (Fig. 3h), in line with reports that *Zbtb46* expression is not exclusive to cDCs (*29*, *54*, *55*).

RORγt^+^ DC showed comparable expression of genes involved in antigen processing and presentation to cDCs and migratory DCs, confirming that RORγt^+^ DCs have the necessary machinery to serve as APC, including expression of *Ly75* and *Wdfy4* (Fig. S3g), both of which can promote cross presentation (*49*, *56*, *57*). RORγt^+^ APCs, including Thetis cells, which also express high levels of *Prdm16, Col17a1,* and *Pigr* have been implicated in promoting Treg differentiation through expression of specific genes, including *Itgb8* (*34*). RORγt^+^ DC did not show an enrichment of Treg promoting genes compared to the other clusters in our analyses and *Itgb8* expression was restricted to the RORγt^+^ eTAC cluster (Fig. 3i, k). Focused multiomic analysis of the RORγt^+^ DC and RORγt^+^ eTAC clusters confirmed that RORγt^+^ eTACs form a separate cluster from RORγt^+^ DCs in ATAC and mRNA profiles and showed that RORγt^+^ DCs can further be divided into 3 clusters (Fig. 3j, k, Fig. S3h). RORγt^+^ DC cluster 2 showed lower expression of *Rorc* than the other two clusters (Fig. 3k), raising the possibility that RORγt^+^ DCs can lose RORγt expression. In this case, cells with *RORγt^Cre^* expression history should outnumber those with active RORγt expression. Indeed, in spleens from two-week-old and adult *RORγt^Cre^Rosa^YFP^RORγt^GFP-knockin^*reporter mice cells with RORγt expression history (YFP) outnumbered those with active RORγt expression (GFP; Fig. 3l).

cDCs from neonates and adults differ functionally and transcriptionally to meet age-specific immune challenges, owing to different environmental signals that transcriptionally imprint these cells across age (*11*, *42*, *58*). Because cDC2 clusters 1 and 2 showed uniform chromatin accessibility but split into two clusters based on gene expression (Fig. S3a, b, d), we first assessed contribution of age to the individual clusters. We found that cells from different ages distributed evenly across ATAC and RNA clusters, with the exception of cDC2 cluster 2 (cDC2_2), which was dominated by cells from the adult time point (Fig. S3i). These data indicate that cDC2 chromatin identity is conserved but environmental signals may shape the cDC2 transcriptional landscape across age. To determine if our experimental set up allowed us to assess differences in gene expression caused by age, we compared cells belonging to the cDC2 metacluster between time points to mimic prior work comparing gene expression in cDC2 across age using bulk RNA sequencing (*11*). This analysis identified 687 differentially expressed genes (Table 3) and showed cDC2 from adult mice were enriched for genes downstream of IFN-γ, tumor necrosis factor-α (TNF-α), IL-2, and IFN-α signaling (Fig. S3j), corroborating our previous findings, and validating the suitability of our data set to compare populations across age. Contrary to cDC2, RORγt^+^ DC clusters showed equal contribution of cells from both ages and only 29 genes were differentially expressed between RORγt^+^ DCs across age (Table 3). These data suggest that RORγt^+^ DCs are transcriptionally stable between 2 weeks of age and adulthood.

Finally, combining chromatin accessibility and gene expression profiles from individual cells we used SCENIC+ (*59*) to identify cell type specific enhancer-driven gene regulatory networks (eRegulons). Dimensionality reduction based on target gene and target region enrichment scores of eRegulons separated the same main cell states as identified above (Fig. 3m). Of note, RORγt^+^ DC and RORγt^+^ eTACs were nearly overlapping in this analysis suggesting that they are closely related cell states. SCENIC+ identified well-known master regulators of cDC1 (Irf8), cDC2 (Runx3), pDCs (Tcf4/E2-2, SpiB), and ILC3 (Rora, Ikzf1, Rorc), confirming the validity of our approach (Fig. S4a). RORγt^+^ DC and RORγt^+^ eTACS shared certain regulatory networks, including Nfat5 and Relb with migDCs and Rel with cDC2, aligning them with DCs (Fig 3n, Fig. S4a). SCENIC+ predicted RORγt^+^ DCs and RORγt^+^ eTACs to be regulated specifically by *Prdm16*, which is in line with specific expression of this transcription factor in RORγt^+^ DC and RORγt^+^ eTACs (Fig. 3n, Fig. S4a). eRegulons of the nuclear receptors Thrb/Nr1a2, Rorc and Ppara were equally predicted by SCENIC+ to regulate RORγt^+^ DCs (Fig. S4a). PRDM16 is a master regulator of brown adipogenesis that in adipocytes directly controls expression of *Ppara*, a nuclear receptor that binds free fatty acids and is involved in lipid metabolism (*60*, *61*). Visualization of the eRegulons formed by Rorc, Thrb and Prdm16 in RORγt^+^ DC suggests cooperativity between these factors in regulating gene expression in RORγt^+^ DCs (Fig. S4b). Accordingly, downstream target genes of PRDM16 showed higher expression in RORγt^+^ DC and RORγt^+^ eTACs than in other cell types and some of these genes appear co-regulated by PRDM16 and RORC (Fig. S4c). Together, these findings establish the unique transcriptional and epigenetic identity of RORγt^+^ DCs. Although showing strong resemblance to migratory DCs, RORγt^+^ DCs are transcriptionally clearly delineated from other DC subtypes and ILC3s. The close epigenetic similarity of RORγt^+^ eTACs to RORγt^+^ DC suggest that AIRE may mark a transcriptional state of RORγt^+^ DCs.

### Identification of RORγt^+^ DCs in human spleen

Comparative transcriptomics are powerful to demonstrate cross species correspondence of APC subtypes (*1*, *12*, *16*). To investigate if RORγt^+^ DCs also exist in human spleen we first turned to a publicly available scRNA-seq dataset of human spleen cDCs enriched as lineage marker negative CD14^-^CD11c^+^HLA-DR^+^ cells from a 61-year-old patient (*16*). We identified 12 clusters, 11 of which corresponded to the previously identified clusters, including cDC1, CLEC10A^-^ cDC2A and CLEC10A^+^ cDC2B, DC progenitor like cells (pre-DC/AS DCs), CCR7^+^ migratory DCs (CCR7^+^ DCs) and one cluster of dividing cells (Fig. 4a and Fig. S5a-c). The 12^th^ cluster was distinguished by *RORC* and *PRDM16* expression (Fig. 4a, b) but lacked expression of the typical ILC3 genes *RORA* and *IL7R*. These cells corresponded to a cluster excluded from the original analyses as putative ILC3 contamination and accordingly scored higher for expression of ILC3 signature genes than the other clusters (Fig. 4c). In murine spleen we identified a set of genes that reliably distinguished RORγt^+^ DCs from cDCs and ILC3s in multiome and scRNA-seq analyses (*11*) (Table 1). Importantly, the *RORC*^+^*PRDM16^+^* cluster in human spleen scored high for these genes (Fig. 4c, Table 1), suggesting that this cluster may correspond to RORγt^+^ DCs identified in murine spleen.

**Figure 4.**
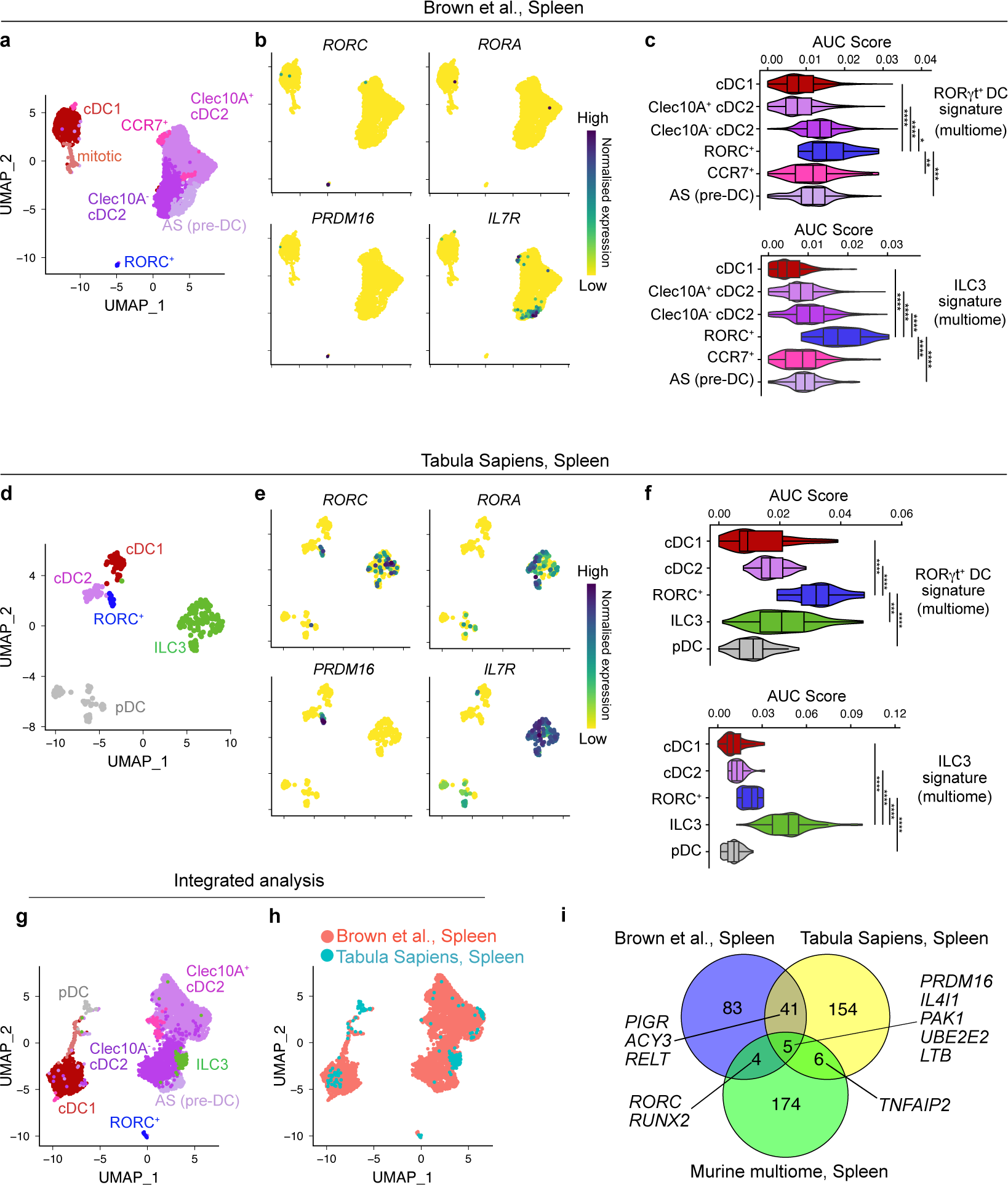
RORγt^+^ DCs exist in human spleen. **a-g.** Annotated UMAP of 4717 cells from scRNA-seq dataset of human splenic DCs. **b.** UMAP feature plots showing *RORC, RORA, PRDM16* and *IL7R*. **c.** Enrichment score for RORγt^+^ DC signature (upper plot) or ILC3 signature (lower plot) from murine multiome dataset calculated for each cluster computed with AUCell package. **d.** Annotated UMAP of 262 cells from scRNA-seq dataset of human spleen. **e**. UMAP feature plots showing *RORC, RORA, PRDM16* and *IL7R.* **f.** Enrichment score for RORγt^+^ DC signature (upper plot) or ILC3 signature (lower plot) from murine multiome dataset calculated for each cluster computed with AUCell package. **g.** Annotated UMAP of integrated scRNA-seq datasets of Brown et al., 2019 and Tabula Sapiens Consortium, 2022, generated using Seurat Integration of scRNA-seq datasets pipeline. **h.** UMAP display of integrated scRNA-seq dataset colored to show contribution of individual datasets to clusters. **i.** Venn diagram showing the overlap of genes distinguishing RORγt^+^ DCs in the indicated mouse and human scRNA-seq datasets. *p (<0.05), **p (<0.01) ***p (<0.001), ****p < 0.0001 **(c, f)** Statistical analysis was performed using Wilcoxon non-parametric ranked sum test to calculate differences in AUCscores between indicated clusters.

To corroborate this further we turned to the spleen dataset from the Tabula Sapiens human reference atlas (*62*). Because RORγt^+^ DCs are scarce, we pre-selected cells annotated as CD1c^+^and CD141^+^ myeloid DCs, pDCs and ILCs, which segregated into 8 clusters that could be categorized as cDC1 (cluster 2), cDC2 (cluster 4), ILC3 (cluster 1, 0, 5), and pDCs (clusters 6 and 3) (Fig. S5d-e). Cells in cluster 7 expressed *RORC* and *PRDM16* but lacked *RORA* and *IL7R*. Importantly, this *RORC^+^* cluster scored highest for genes distinguishing RORγt^+^ DCs in mouse spleen, whereas ILC3s showed the highest enrichment of ILC3 signature genes (Fig. 4f, Table 1). This *RORC^+^*cluster contained cells from multiple donors aged 59, 61, and 69 years and did not segregate by sequencing method or cell cycle phase, validating it as a bona fide population (Fig. S5f). Finally, we integrated both datasets and found that *RORC^+^* cells from both data sets clustered together and away from DCs and ILCs (Fig. 4g, h). Comparative gene expression analyses in the mouse and human spleen data sets identified *PRDM16, UBE2E2, IL4I1, PIGR* and *LTB* as markers that distinguish the *RORC^+^* clusters in human and mouse spleen (Fig. 4i). Although scRNA-seq analyses does not allow to distinguish the RORγt isoform of RORC, collectively, these cross-study and cross-species analyses demonstrate that RORγt^+^ DCs that we identified in mouse spleen also exist in human spleen.

### RORγt^+^ DCs exist across lymphoid and non-lymphoid tissues in mice and humans

Janus cells/RORγt^+^ eTACs and Thetis cells are found at barrier associated lymphoid tissues (*33*, *34*, *38*) and in humans RORγt^+^ APCs with characteristics of both cDC2 and ILC3s have also been described in tonsils and nasal associated tissue (*41*). We therefore asked if RORγt^+^ DCs exist in organs other than spleen. We first profiled mLN and small intestinal lamina propria (siLP) from *RORγt^GFP^* mice by flow cytometry. Migratory cDCs in non-lymphoid tissues express CD127 (*29*, *63*) (Fig. S6a), leading us to gate cDCs irrespective of this marker as CD90^-^CD11c^+^MHCII^+^ cells lacking the monocyte marker CD64. Within this gate we detected RORγt^+^ DCs at all ages examined and found their frequency was highest in neonatal mLN and declined with age (Fig. 5 a,b). Although the absolute number of cells recovered from neonatal mLN varied, we consistently detected the highest number of RORγt^+^ DCs at two weeks of age (Fig. 5a,b, Fig. S6b). In siLP we observed a similar trend. The frequency of RORγt^+^ DCs within CD11c^+^MHCII^+^ cDCs was highest in the neonatal period and absolute numbers peaked at two weeks of age. However, it needs to be taken into account that for technical reasons Peyer’s patches, which also contain RORγt^+^ APCs resembling cDCs (*42*) were removed from SI only for mice older than two weeks of age (Fig. 5b). Phenotypically, RORγt^+^ DCs were distinct from MHCII^+^ILC3s (CD90^+^CD127^+^MHCII^+^GFP^+^ cells, Fig. S6a, b,e,f). In mLN and siLP from one-week-old, but not adult mice, RORγt^+^ DCs expressed CD11b (Fig. S6 c, f). In both tissues RORγt^+^ DCs expressed high levels of MHCII, consistent with a migratory DC phenotype and lacked the cDC1 marker XCR1, although some RORγt^+^ DCs expressed XCR1 in adult mLN (Fig. S6c,f). RORγt^+^ DCs also lacked the ILC3 marker CXCR6 in mLN (Fig. S6d). In the lung, RORγt^+^ DCs could also be detected across age but were generally least frequent in this organ (Fig. 5c, Fig. S7a). Nonetheless, the numbers of RORγt^+^ DCs peaked at two weeks of age (Fig. 5c). Pulmonary RORγt^+^ DCs lacked XCR1 but notably expressed CD11b in neonatal and adult mice (Fig. S7b) and resembled cDC1 and cDC2 in terms of CD11c and MHCII expression, while ILC3s exhibited lower CD11c and MHCII expression (Fig. S7b).

**Figure 5.**
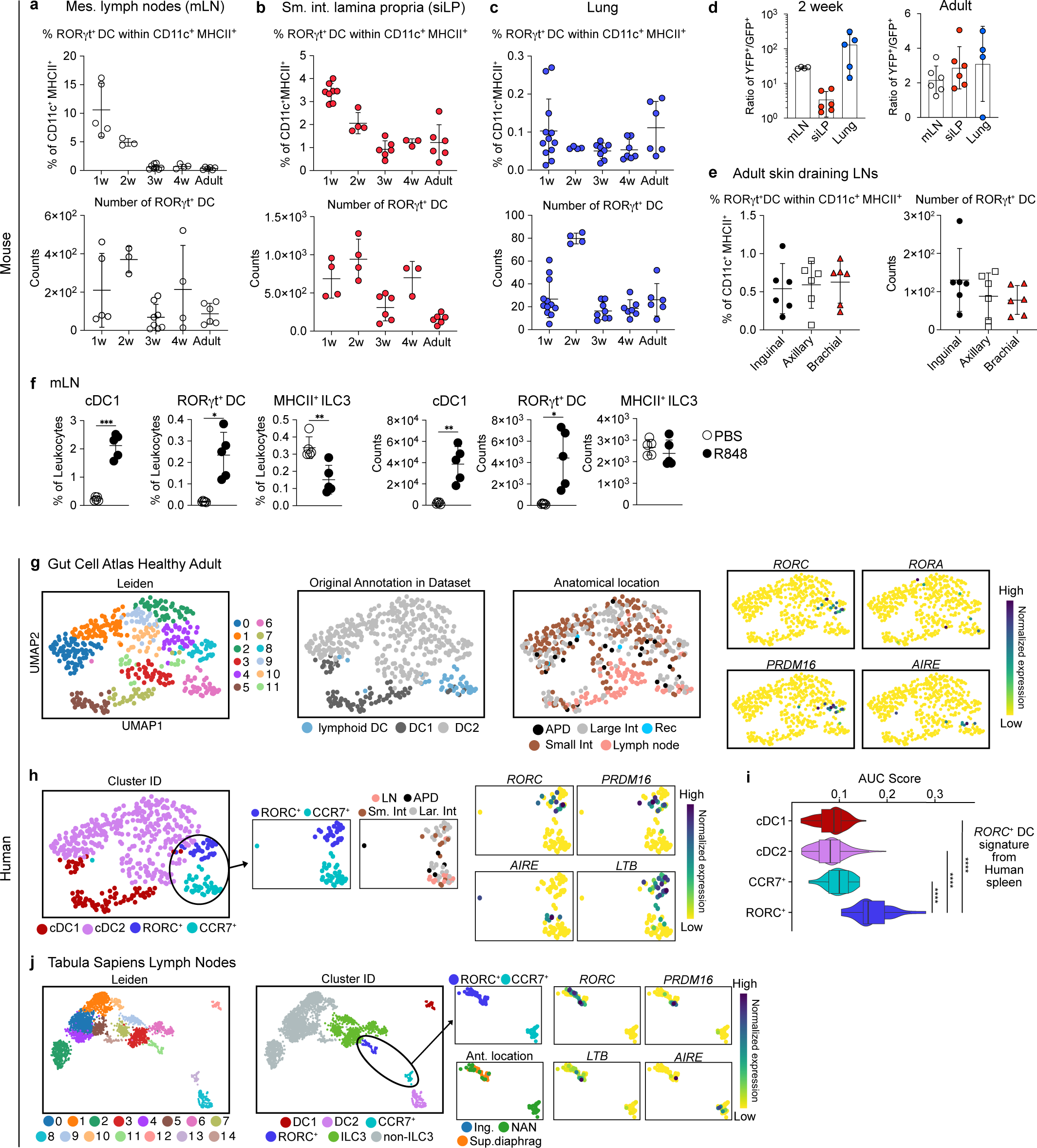
RORγt^+^ DCs exist across lymphoid and non-lymphoid tissues in mice and humans. **a-c.** RORγt^+^ DCs were quantified in mLN **(a)**, siLP **(b)** and lung **(c)** of *RORγt^GFP^* reporter mice (gating see in Fig. S6 and S7a). **a.** Frequency of RORγt^+^ cells within CD11c^+^MHCII^+^ cells (upper plot) and number of RORγt^+^ cells (lower plot) in mLN of mice of indicated ages (1 week n=5; 2 week n=3; 3 week n=8, 4 week n=4; Adult n=6). **b, c.** Frequency of RORγt^+^ cells within CD11c^+^MHCII^+^CD64^-^ cells (upper plot) and number (lower plot) in siLP (1-week-old n=4-8; 2-week-old n=4; 3-week-old n=6, 4-week-old n=3; Adult n=6) and lung (1-week-old n=11; 2-week-old n=4; 3-week-old n=7, 4-week-old n=6; Adult n=6) from mice of indicated ages. **a-c**. **d.** mLN, siLP and lung from *RORγt*^Cre^*Rosa*^YFP^*RORγt*^GFP/wt^ mice were analyzed by flow cytometry as in Fig. S7d. The ratio CD11c^+^MHCII^+^YFP^+^ cells (RORγt expression history) over CD11c^+^MHCII^+^GFP^+^ cells (active RORγt expression) is plotted. Each dot represents one mouse. **e.** RORγt^+^ DC from axial, brachial and inguinal skin draining LN were gated as in Fig. S7e and quantified. The frequency of RORγt^+^ cells within CD11c^+^MHCII^+^ cells (left) and number of RORγt^+^ DCs (right) is shown. **f.** Quantification of cDC1, MHCII^+^ILC3 and RORγt^+^ DCs in mLN of 11-day old *RORγt^GFP^* reporter mice 24 hours after oral administration of R848. **g-j.** 480 cells annotated as lymphoid DCs, DC1 and DC2 from healthy adult donors from the human gut cell atlas were reclustered using *Scanpy* into 12 clusters. UMAPs indicating cluster distribution, cell annotation and anatomical location are shown. Right plots show expression of *RORC*, *PRDM16*, *RORA* and *AIRE*. **h.** UMAP from **(g)** annotated using signature genes (Fig. S8a) and zoomed in display of *RORC^+^*and *CCR7^+^* clusters to illustrate anatomical location and expression of *RORC*, *PRDM16*, *AIRE* and *LTB* in individual cells. **i.** Enrichment score for genes delineating *RORC*^+^ cells in human spleen computed for each cluster with AUCell package. **j**. 2668 cells annotated as CD1c^+^DC, CD141^+^ DC, cDC, HSC, ILC and pDC in the Tabula Sapiens lymph node reference set were reclustered into 15 clusters. UMAP displaying Leiden clustering and annotation based on signature genes (Fig. S8c). Zoomed in display of *CCR7^+^* and *RORC^+^* clusters to visualize anatomical location and expression of the indicated genes. Statistical analysis in **(i)** was performed using Wilcoxon non-parametric ranked sum test to calculate differences in AUCscores between indicated clusters. *p (<0.05), **p (<0.01) ***p (<0.001), ****p < 0.0001. Data in **a-f** are pooled from at least 2 independent experiments per time point, each dot represents a biological replicate.

Similar to our observations in spleen, we found that cells with RORγt expression history outnumbered those with active RORγt expression in lung, mLN and siLP of adult mice by about 3 times. In lung and mLN from two-week-old mice, respectively, cells with *RORγt^Cre^* expression history were even 100 and 10-fold more frequent than those actively expressing RORγt (Fig. 5e, Fig. S7d). These data suggest that some RORγt^+^ DCs may downregulate RORγt in response to tissue specific signals. RORγt^+^ DC from spleen transcriptionally resembled Janus cells/RORγt^+^ eTACs that have primarily been described in lymph nodes draining non-mucosal organs (*38*). Importantly, we also detected RORγt^+^ DCs in skin draining lymph nodes which expressed high levels of MHCII, again consistent with a migratory DC phenotype (Fig. 5f, Fig. S7e,f).

To address if RORγt^+^ DCs have the capacity to migrate from intestine to mLN, we gavaged neonatal mice with R848, an inflammatory stimulus that induces cDC migration to mLN (*64*). Indeed, 24 hours after R848 gavage we observed an increase in the frequency and number of cDC1, but not ILC3s, in mLN, which served as positive and negative controls for migration, respectively (Fig. 5f). RORγt^+^ DCs also increased in frequency and number in mLN (Fig. 5f), suggesting that oral R848 stimulates RORγt^+^ DCs to migrate to mLN. Of note, RORγt^+^ APCs resembling cDCs in neonatal Peyer’s patch also become activated upon R848 stimulation (*42*). Thus, RORγt^+^ DCs phenotypically distinct from ILCs and other cDC subtypes exist in mLN, siLP and lung, and bear classical migratory features of cDCs upon inflammatory triggers, validating their affiliation with DCs.

To address if RORγt^+^ DCs exist in non-lymphoid human tissues, we turned to a single-cell atlas encompassing second trimester to adult intestine and mLN (*65*). We first selected cells from adult healthy donors ranging from 25-75 years of age, annotated in the original data set as DC1, DC2 and lymphoid DCs. Leiden clustering of these cells revealed 12 clusters, of which cluster 8 showed expression of *RORC, PRDM16* and *LTB,* but not *RORA,* and mainly originated from small and large intestine (Fig. 5g,h). The other clusters could be broadly categorized into cDC1, cDC2, and CCR7^+^ DCs based on signature genes (Fig. S8a). Of note, *AIRE* expression was detected in cluster 6, which was classified as CCR7^+^ migratory DCs based on CCR7 expression and the original annotation as lymphoid DC (Fig. 5g, h, Fig. S8a). The *RORC^+^* cluster scored higher for expression of genes that distinguished RORγt^+^ DCs in human spleen (Fig. 5i and Table 5). Importantly, the *RORC*^+^ cluster contained cells derived from small and large intestine of multiple donors (Fig. 5g, Fig. S8a), establishing that RORγt^+^ DCs exist in human intestine. Importantly, *RORC* and *PRDM16* expressing cells were also identified in small intestine of healthy pediatric donors and pediatric Crohn’s disease patients ranging from 4-14 years of age (Fig. S8b), confirming that RORγt^+^ DCs are conserved in human intestine across age. *RORC, PRDM16* and *LTB* expressing cells were also detected in the Tabula Sapiens reference data set for lymph nodes (*62*) (Fig. 5j, Fig. S8c,d). These cells clustered distinctly from ILCs and DCs and expressed *CCR7*, suggesting that they may have migrated from the periphery to the draining lymph node (Fig. 5j). Of note, some of the cells within the *RORC*^+^ cluster in lymph nodes also expressed *AIRE* (Fig. 5j). Cells within this cluster were again derived from multiple donors and originated from subdiaphragmatic LN and from LN with unspecified anatomical origin (Fig. S8c). Collectively, these analyses show that cells transcriptionally resembling RORγt^+^ DC can be identified in various human tissues.

### Integrative transcriptional analyses align RORγt^+^ DCs and other RORγt^+^ APCs across mouse and human tissues

Finally, we aimed to more comprehensively address if RORγt^+^ DCs have conserved transcriptional features that distinguish them from other APCs across tissues. First, we performed comparative gene expression analyses of RORγt^+^ DCs relative to other cells in the mouse and human data sets from spleen, lymph nodes and intestines described above. We also included a scRNA-Seq dataset of CD11c^+^MHCII^+^ cells from murine neonatal SI, in which we could transcriptionally identify a cluster of RORγt^+^ DCs to these comparative analyses (Fig. S9a). By identifying commonalities, we revealed a set of genes that reliably distinguished RORγt^+^ DCs across all of the above datasets (Tables 1, 5, Fig. 6).

**Figure 6.**
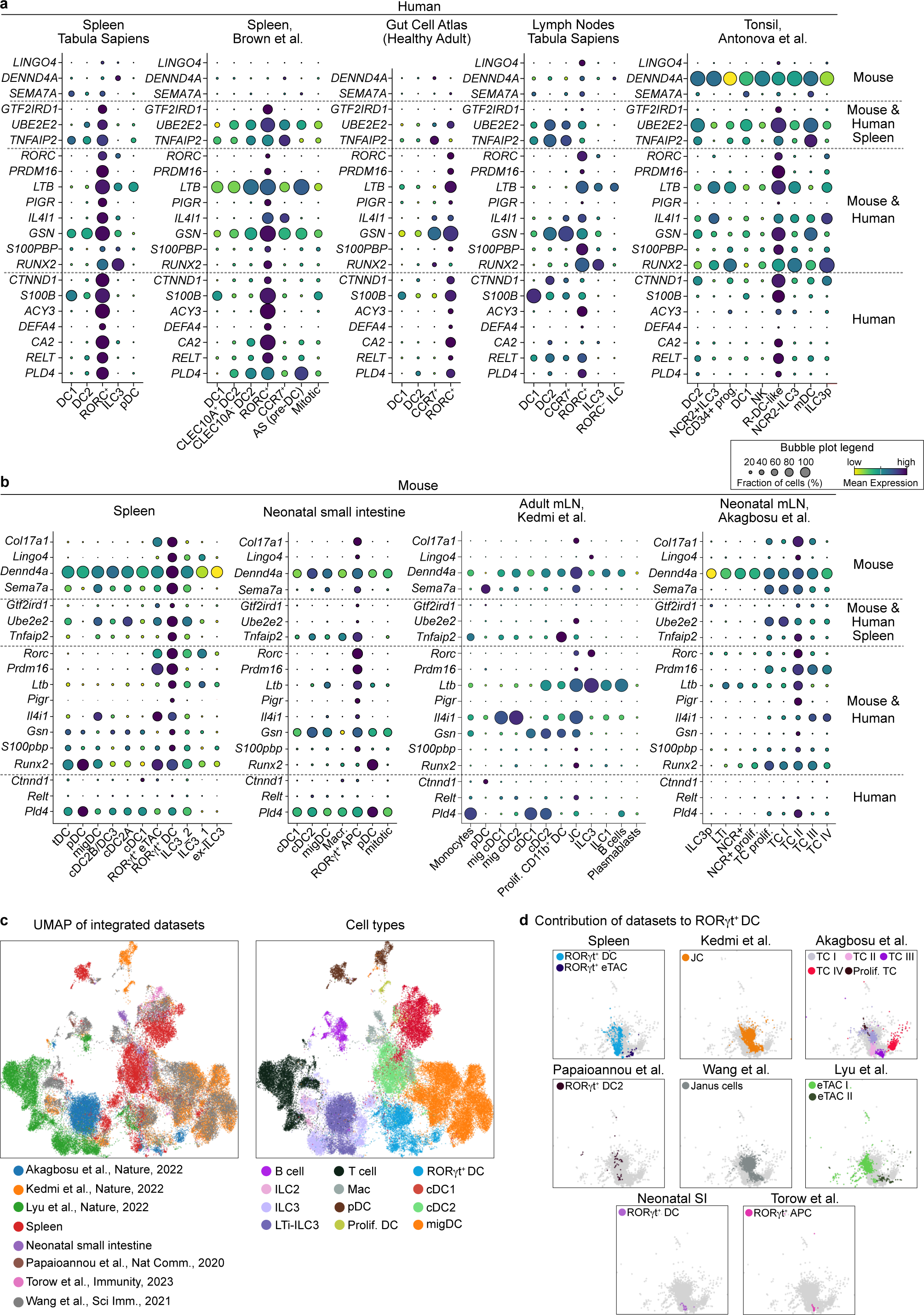
Integrative transcriptional analyses align RORγt^+^ DCs and other RORγt^+^ APCs across mouse and human tissues. **a, b.** Bubble plots showing expression of selected genes deduced from comparative genes expression analyses that distinguish RORγt^+^ DCs in various human **(a)** and murine **(b)** scRNA-seq data sets. Genes are ordered according to the species and organs they were found in to differentiate RORγt^+^ DCs from other cell types. **c.** The indicated murine data sets were integrated. UMAP display of the integrated datasets colored by data set and annotated cell type to show contribution of the individual datasets to each cluster. **d.** Zoomed in display of the RORγt^+^ DC cluster from the UMAP in **(c).** Colored in grey are all cells contributing to the cluster. In each panel the contribution of the specified populations from the indicated publications is shown.

We then checked the expression of these genes in various human and murine datasets. We found that ‘R-DC-like cells’ from tonsil (*41*) showed higher expression of these genes compared to other DC and ILC3 populations in the same data set suggesting these cells are homologous to RORγt^+^ DC described in our study (Fig. 6a). In murine data sets we found that RORγt^+^ APCs from neonatal Peyer’s patch (*42*), Thetis cells from neonatal mLN (*34*), Janus cells from mLN (*33*) and sdLN (*38*), and RORγt^+^ eTACs from mLN (*27*) all showed higher expression of these genes compared to other populations within the respective datasets (Fig. 6b, Fig. S10a). Of note, TCII showed highest expression of RORγt^+^ DC signature genes, while TCIV showed lowest expression of these genes (Fig 6a). In line with previous analyses (*34*, *40*), TCIII and TCIV showed high expression of the eTAC gene *Tnrfsf11a*, although Aire expression was only observed in TCI and TCII (Fig. S9c).

These above data indicate a strong transcriptional similarity between various subtypes of RORγt-expressing APCs described in different tissues and different stages of life to RORγt^+^ DCs identified in our study, raising the possibility that they all belong to the same lineage of cells. To address this, we performed data integration of the above mentioned published murine datasets together with our own datasets from murine spleen in adults and neonates (Fig. 3) (*11*) and neonatal small intestine (Fig. S9a). After unsupervised dataset integration using scVI (*66*) we could see that cells from individual datasets were distributed evenly across Leiden clusters (Fig. 6c and Fig. S10b). Applying the original cell type annotations revealed that cells annotated as the same broad cell types in the individual datasets (cDC1, cDC2, migDCs, pDCs, Mac, T cells, B cells, ILC3) clustered together in the integrated analysis irrespective of study and tissue of origin and that the Leiden clusters corresponded well to the different cell types included (Fig. 6c, Fig. S10c). Importantly, we found that all cells annotated as eTACs, Janus cells, Thetis cells, or RORγt^+^ APCs, in their original datasets clustered together with RORγt^+^ DC identified in our study (Fig. 6d) into a transcriptionally distinct cluster that we therefore termed RORγt^+^ DC (Fig. 6c). Of note, although TCIV fell within the RORγt^+^ DC cluster, TCIV were somewhat spatially separated and did not notably overlap with cells from other datasets (Fig. 6d). This comprehensive meta-analysis unequivocally establishes RORγt^+^ DC as a transcriptionally distinct lineage of immune cells that is conserved across tissues and species, and encompasses previously described populations including RORγt^+^ eTACs, Janus and Thetis cells (Fig. S10d).

## DISCUSSION

The delineation of cDCs from other APCs has historically been blurry (*1*, *2*, *67*) but with the use of genetic tools it has become clear that the functional specialization of cDC subtypes is critical for successful immunity (*1–3*). Here, we define RORγt^+^ DCs as bona fide APCs conserved across tissues and species with striking similarities to cDCs, including ability to migrate to lymph nodes and activate naïve T cells. RORγt^+^ DCs possess a unique transcriptional identity that aligns them with other recently described APC populations, including R-DC-like cells in humans (*41*), Janus cells/eTACs in lymph nodes (*33*, *38*), Thetis cells in neonatal mLN (*34*) and RORγt^+^ APCs in neonatal Peyer’s patches (*42*), suggesting that these cells all belong to the same lineage of APC (Fig. S10d). Interestingly, TCIV did not show a direct overlap with cells from other data sets in our integration analysis, raising the possibility that these cells may be a specific transcriptional state of RORγt^+^ DCs or restricted to neonatal mLN (*34*).

While RORγt^+^ APCs have primarily been linked to T cell tolerance and induction of Tregs, RORγt^+^ DCs from spleen express genes involved in antigen processing and presentation and lack a particular tolerogenic profile. Accordingly, we find that RORγt^+^ DCs from spleen, like cDCs, respond to inflammation with activation of naïve T cells (*41*). Similarly, R-DC-like cells in humans can stimulate allogeneic CD4^+^ T cells, although their APC potential has not been compared to that of cDCs (*41*). Further, we show that RORγt^+^ DCs migrate to mLN following an intestinal inflammatory trigger. These data suggest that RORγt^+^ DC can serve as bona fide immune sentinels. Tolerogenic immune functions in RORγt^+^ APCs have been linked to AIRE expression (*33*, *38*) and AIRE expression is restricted to subtypes of RORγt^+^ DCs in our and previous studies (*33*, *34*). In tumor associated macrophages AIRE marks a distinct transcriptional cell state (*68*) and AIRE drives functional programs in DCs and ILC3s (*32*, *52*). Since RANK signaling has been shown to induce AIRE in eTACs and ILC3s (*37*, *38*) it is possible that RANKL or other tissue or age-specific signals shape AIRE expression or other transcriptional and functional attributes of RORγt^+^ DCs. It therefore follows to conclude that RORγt^+^ DC are versatile antigen presenting cells that can drive inflammatory and anti-inflammatory immunity in a context-dependent manner. As such, it will be important to define the tissue specific signals regulating this cell type. Owing to their expression of CD11c, ZBTB46, and RORγt, RORγt^+^ DCs are important to consider in the interpretation of studies using promoters for the above genes to drive Cre or other transgenes, such as diphtheria toxin receptor, to define immune functions of cDCs or RORγt^+^ APCs.

We provide a flow-based strategy by which RORγt^+^ DCs can accurately be identified across tissues either by using *RORγt^GFP^*reporter mice or intracellular staining using an anti-RORC antibody. However, their limiting numbers and phenotypic overlap with the above-mentioned populations makes it difficult to probe their functions. Thus, models to specifically manipulate this cell type are urgently needed to understand its functions in immunity. In the absence of such models, the antigen-targeting approach used here will prove useful to determine the specific functions of RORγt^+^ DCs compared to other APC subtypes in the context of different adjuvants. It will be important to determine if the putative transcriptional regulators for RORγt^+^ DCs identified in our study, such as *Prdm16* and *Thrb/Nr1a2*, specifically drive the development of RORγt^+^ DCs. Of note, SCENIC+ predicted RORγt itself to drive transcriptional programs in RORγt^+^ DCs but it is unclear if these cells require RORγt for their development. Such investigations require specific genetic models to trace expression of RORγt even after genetic ablation or alternative markers to specifically identify this cell type, which we have not yet been able to confirm. To overcome this, we have also identified a set of genes, including PRDM16, that reliably defines RORγt^+^ DCs across tissues and species in single cell based transcriptional profiling.

Importantly, RORγt lineage tracing experiments show that some cells that have previously expressed RORγt downregulate it. Thus, RORγt itself may not be a good lineage defining marker of RORγt^+^ DCs. Specifically in neonatal lung where DC-like cells with *RORγt^Cre^* expression history vastly outnumber cells with active RORγt expression. This is interesting as neonatal lung is considered an environment that promotes type 2 immunity (*69*), while RORγt is generally associated with type 3 immunity. Thus, factors specifically expressed in neonatal lung may suppress RORγt expression, highlighting the need to define the tissue specific signals regulating this cell type. RNA velocity analyses in humans suggest that RORγt^+^ DCs may serve as alternative progenitors for cDC or ILC3-like cells (*41*). Although we found no evidence for that in steady state neonatal spleen (*11*) it is noteworthy that PRDM16, the putative transcriptional regulator of RORγt^+^ DCs, is a known regulator of stem cell quiescence (*70*). Thus, the progenitor potential of RORγt^+^ DCs requires future investigation, especially in situations of inflammation.

Future work is needed to better define the ontogenetic relationships of RORγt^+^ DCs to other immune cells. We demonstrate that RORγt^+^ DCs are ontogenetically distinct from ILC3 and do not arise from Clec9a-expressing cDC-restricted progenitors but their specific progenitor remains to be identified. The fact that RORγt^+^ DCs are transcriptionally conserved across tissues and age highlights that these cells are not a “state” solely acquired at a specific tissue site. Therefore, we propose to term these cells RORγt^+^ DCs to acknowledge their uncertain origin while abiding to their phenotypic, functional, and transcriptional resemblance to dendritic cells (*1*, *35*, *41*, *47*). Our functional characterization coupled to robust meta-analysis tying RORγt^+^ DCs to previously published populations of RORγt^+^ APCs show that RORγt^+^ DCs are bona fide APCs with a versatile functional spectrum ranging from inducing T cell activation to tolerance. This versatility makes them an attractive target for therapeutic manipulation.

## MATERIALS AND METHODS

### Mice

Tg(*Rorc*-EGFP)1Ebe (*71*), C57BL/6-*Flt3l*^tm1Imx/TacMmjax^ (MMRRC Stock No:37395-JAX), OTII mice (Tg(TcraTcrb)425Cbn; Jackson Laboratory Stock No: 004194) crossed to a Thy1.1 (CD90.1) background, *Clec9a*^tm2.1(icre)Crs^ (Jackson Laboratory Stock No: 025523), Gt(ROSA)26Sor^tm9(CAG-tdTomato)Hze^ (*Rosa26*^lox-STOP-lox-tdtomato^) (Jackson Laboratory Stock No: 007909), and C57BL/6J RccHsd were bred and maintained at the Biomedical Center, LMU, Munich. *Rag2^-/-^Il2rg^-/-^, Rorc(*γ*t)^GFP/wt^* (*72*) crossed to *Rorc(*γ*t)*^cre^(*43*) and *Rosa26*^YFP^ mice were bred and maintained at the Federal Institute for Risk Assessment (Berlin, Germany) and the Research Institute for Experimental Medicine (FEM) of the Charité (Berlin, Germany). *RORγt*^GFP^*RORγt*^Cre^*Rosa26*^RFP^ were a gift from Vasileios Bekiaris and bred and maintained at the Technical University of Denmark BioFacility. All mice were housed under specific pathogen free conditions, with a 12 h dark/light cycle, in individually vented cages. Food and water were provided *ad libitum*. Both female and male mice were used in this study. Where applicable age and sex-matched mice were used for experiments. In most experiments littermates were used. All animal procedures were performed in accordance with national and institutional guidelines for animal welfare and approved by the Regierung of Oberbayern, Landesamt für Gesundheit und Soziales or the Danish Animal Experiments Inspectorate.

### Cell isolation for flow cytometry

Spleens were cut into small pieces and enzymatically digested in 1mL RPMI containing 200 U/mL Collagenase IV (Worthington) and 0.2 mg/mL DNaseI (Roche, 11284932001) for 30 minutes at 37°C while shaking at 180 rpm. Digested suspensions were passed through a 70 μm strainer and washed with FACS buffer (PBS, 1% fetal calf serum (FCS), 2.5 mM EDTA, 0.02% sodium azide). Erythrocytes were then lysed with Ammonium-chloride-potassium (ACK) lysis buffer (1mM EDTA, 1.55 M NH_4_Cl, 100mM KHCO_3,_ filtered) for two minutes at room temperature (RT) followed by a wash with FACS buffer. Cell pellets were resuspended in FACS buffer for further analysis.

Mesenteric lymph nodes (mLN) from 2-3 one week old mice were pooled prior to processing. For all other timepoints mLN from individual mic were processed. Tissue was processed as above but without erythrocyte lysis. Axillary, brachial, and inguinal skin draining lymph nodes from adult mice were enzymatically digested in 1mL of RPMI with 200 U/mL Collagenase IV (Worthington) and 0.2 mg/mL DNaseI (Roche) for 30 minutes at 37 °C while shaking at 400 rpm. Digested samples were passed through 70 μm strainer, washed and resuspended in FACS buffer for further analysis.

Small intestines (SI) were isolated and flushed with ice-cold wash medium (Hank’s Balanced Salt solution (HBSS), 5 mM EDTA, 5% FCS, 10mM HEPES, without calcium or magnesium). For mice 3 weeks or older, Peyer’s patches were then removed. Intestines from mice younger than 3 weeks were cut open with a scalpel and washed in ice cold PBS. SI from one-day old mice were pooled by 2-3. Tissues were cut into pieces and incubated in wash medium and 1 mM dithiothreitol (DTT) for 30 minutes with shaking (180 rpm). Supernatant containing epithelial cells and intraepithelial lymphocytes was discarded. Samples were then washed with digestion medium (HBSS with Calcium and Magnesium, 5% FCS and 10mM HEPES) and then cut to smaller pieces for enzymatic digestion. Samples were then digested in 3mL digestion medium containing 400 U/mL Collagenase IV (Worthington) and 0.4 mg/mL DNaseI (Roche, 11284932001), for 30 minutes at 37 °C with shaking (180 rpm). Digested samples were passed through 100 μm strainer and washed with gut medium (RPMI with 1% Glutamine, 1% Penicillin/Streptomycin, 10% FCS, 10 mM HEPES, and 50 μM ϕ3-Mercaptoethanol). Supernatant was removed by suction with vacuum pump. Leukocytes were enriched by 70%-37%-30% Percoll (Sigma Aldrich) gradient centrifugation (2000 rpm, RT, 30 minutes, Accel=3, Decel=0).). Cells were collected from the 70%-37% interphase and washed once in gut medium and resuspended in FACS buffer (without sodium azide) for further analyses.

For lung isolation mice were perfused with ice cold PBS. Lungs from mice younger than 3 weeks were isolated without perfusion. Tissue was minced and enzymatically digested in 1mL RPMI containing 200 U/mL Collagenase IV (Worthington) and 0.2 mg/mL DNaseI (Roche) for 1 hour at 37 °C with shaking (180 rpm). Digested tissue was passed through a 70μm strainer and washed with FACS buffer. Leukocytes were enriched by gradient centrifugation as above.

### OTII enrichment

Spleens from OTII mice were collected in 1mL RPMI, mechanically disrupted and passed through a 70 μm strainer and washed with FACS buffer without sodium azide ((PBS, 1% fetal calf serum (FCS), 2.5 mM EDTA). Naïve CD4^+^ OTII cells were isolated using MojoSort™ mouse CD4 T Cell Isolation Kit (BioLegend) according to the manufacturer’s instructions. OTII cells were labelled with CellTrace™ Violet (CellTrace™ Violet Cell Proliferation Kit, Invitrogen) according to manufacturer’s recommendations.

### Flow cytometry

Cells were first incubated with 50μL purified anti-mouse CD16/32/FcBlock for 10 minutes at 4°C in 96-well v-bottom plates. Next, cell surface epitopes were stained for 30 minutes at 4°C by addition of 50μL antibodies in a 2X mastermix (total staining volume of 100μL). Fixable Viability Dye eFluor 780 (Thermo Fisher Scientific) was added at this step. To preserve GFP signal after intranuclear staining, cells were pre-fixed with 2% paraformaldehyde at RT for 15 min. Intracellular staining for cytokines was performed using Intracellular Fixation & Permeabilization Buffer Set and for transcription factors with the FOXP3 Transcription Factor Staining Set (both Thermo Fisher Scientific) according to the manufacturer’s instructions after staining for surface epitopes. Anti-RORC staining was performed for 1hr at RT. Cells were quantified by using CountBright™ Absolute Counting Beads (Thermo Fisher Scientific). Flow cytometry data was collected using LSR Fortessa (BD Biosciences) and analyzed using FlowJo software V10.8.1 (Tree Star). Antibodies used for flow cytometry are provided in Table 6.

### Cell sorting

For sorting of APC populations splenocytes were depleted of T cells (CD3e), B cells (CD19), Neutrophils (Ly6G) and Erythroid cells (Ter119) by staining with FITC-conjugated antibodies followed by anti-FITC magnetic bead (Miltenyi) negative selection with LS columns (Miltenyi) according to manufacturer’s instructions. Cells were stained in total volume of 600μL in 50mL conical tubes. For in vitro co-culture assays PBS + 10% FCS was used as collection medium. Cells for single cell multiome analyses were collected in complete RPMI (1% L-Glutamine, 1% Penicillin/Streptomycin, 10% FCS, 1% Sodium Pyruvate, 0.1% ϕ3-Mercaptoethanol and 1% non-essential amino acids). Cell sorting was performed using BD FACSAria Fusion (BD Biosciences). Antibodies used for sorting are provided in Supplementary Table 1.

### Sorting from neonatal small intestine for scRNA-seq

Single cell suspensions from small intestines from one-day old (n=5, pooled as 2 and 3 for processing) and nine-day old (n=3) *Clec9a*^Cre^ *Rosa*^TOM^ mice were generated. Cells from each timepoint were stained with TotalSeq^TM^ anti-mouse Hashtag antibodies (BioLegend) when staining for cell surface epitopes. Live CD45^+^ singlets, CD19^-^ CD90.2^-^ CD11c^+^ MHCII^+^ cells were sorted from each timepoint and pooled in a 1:1 ratio for sequencing using the 10X Genomics Chromium Next GEM Single Cell 3’ v3.1 (Dual Index) with Feature Barcode Technology for Cell Surface Protein protocol.

### In vitro T cell proliferation

MHCII^+^ ILC3, cDC2, and RORγt^+^ DC were sorted as described above (gating strategy Fig S2). 250 cells from each population were pulsed with 10μg/mL Ovalbumin peptide (OVA_323-339_, Invivogen) in a 96-well v-bottom plate for 3 hours, then washed and resuspended in complete RPMI (cRPMI - 1% L-Glutamine, 1% Penicillin/Streptomycin, 10% FCS, 1% Sodium Pyruvate, 0.1% β-Mercaptoethanol and 1% non-essential amino acids) and co-cultured with 2500 naïve CTV-labelled OTII cells at a 1:10 ratio. Cultures were supplemented with 5 ng/mL TGF-β, 10 μg/mL anti-IL-4, and 10 μg/mL anti-IFN-γ for Treg polarization, and 5ng/mL TGF-β, 20ng/mL IL-6, 10μg/mL anti-IL-4, and 10μg/mL anti-IFN-γ (all Biolegend) for Th17 polarizing condition. Cells were cultured in a total volume of 200μL. After 3.5 days of culture at 37°C, supernatant was collected from all samples, following which cells were restimulated with 10ng/mL phorbol 12-myristate 13-acetate (PMA) (Calbiochem) and 1μg/mL Ionomycin (Sigma-Aldrich) for a total of 5 hours. 2 hours after start of restimulation 5μg/mL of Brefeldin A (BioLegend) was added to each well for the remaining 3 hours. Cytokines and FOXP3 were detected by intranuclear staining as described above.

### In vivo targeting

Adult *RORγt^GFP^* mice were injected intraperitoneally (i.p.) with 10 μg anti-CLEC4A4-OVA or 10μg OVA-coupled-isotype matched control antibody as described (*49*) plus 0.2 μg/g body weight of CpG-B ODN 1826 (InvivoGen). 12 hours later 300 cDC1, cDC2, and RORγt^+^ DCs were sort purified from spleens and co-cultured with 3000 naïve CTV-labelled OTII cells in cRPMI. 3.5 days later OTII proliferation was assessed by flow cytometry.

### R848 treatment

10-day old *RORγt^GFP^* mice were orally gavaged with 2μg R848 (Invivogen) in 50μL PBS or received 50μL PBS as control. 24 hours later organs were isolated and processed as above.

### Single cell multiome sequencing of RORγt^+^ DCs, cDCs and ILC3s

MHCII^+^ ILC3, CD11c^+^MHCII^+^ DC and RORγt^+^ DCs were sorted from of 2-week-old (P13, n=2) or adult (8-week-old, n=3) *RORγt^GFP^Clec9a*^cre/wt^*Rosa26*^Tom/wt^ mice into cRPMI as described above. EDTA-free buffer was used in all steps for processing of spleen tissue. For each timepoint, cells were pooled in a ratio of approx. 1:1:10 (RORγt^+^ DCs: MHCII^+^ ILC3: CD11c^+^ MHCII^+^ DC) and nuclei were isolated following the 10X Genomics Low Cell Input Nuclei Isolation protocol. Briefly, 20,000 cells of mixture were lysed for 3 minutes on ice and resuspended in diluted Nuclei Buffer. Lysis efficiency and nucleus quality were assessed after trypan blue staining by microscopy. 8400 nuclei for the adult timepoint and 10,400 nuclei for the 2 week timepoint were loaded for transposition. Gene expression and ATACseq libraries were prepared with the 10X Genomics Chromium Next GEM Single Cell Multiome ATAC + Gene Expression Kit according to manufacturer’s instructions. Concentration and purity of the libraries were analyzed at the required steps with a Aligent TapeStation. Libraries from each timepoint were pooled and sequenced together in accordance with 10X Genomics recommended sequencing depth, using NextSeq1000/2000.

### Multiome computational analysis

Alignment: *Cell Ranger ARC v2.0.2* was used to map the multiome reads against the mm10 reference genome customized to include sequences for *eGFP*, and *tomato*.

Gene expression analyses: The raw reads of the 2-week and the adult samples were combined and genes that were present in fewer than 5 cells were removed. Cells were included if they had between 500 and 5,000 genes detected, fewer than 30,000 counts and less than 1% mitochondrial reads. Library size normalization was done using *scran* before log-transforming the counts. Cell cycle analysis was done in *Scanpy.* Initial Leiden clustering was performed at resolution of 0.5 and expression of marker genes and scoring based on published signatures for cell types (Table 1) was used to annotate the clusters. Based on marker gene expression, cluster 7 was re-clustered at resolution of 0.05 to separate migDCs from eTACs and cluster 3 was re-clustered at resolution of 0.1 to separate tDCs from cDC2s. Clusters were compared between RNA and ATAC to focus on the cellular subtypes that were identifiable from both omics. Differential expression analysis was performed between groups of interest using *Scanpy’s sc.tl.rank_genes_groups* on all genes. Significantly differentially expressed genes were defined by having an adjusted p-value below 0.05 and an absolute log_2_-fold change greater than 0.5. Gene set enrichment analysis was done using *gseapy* (*73*).

ATAC analyses: The bam files output by Cell Ranger ARC were merged for the 2-week and adult sample and peaks were called using *macs3.* Count matrices were built for the 2-week and adult sample using *epiScanpy’s epi.ct.peak_mtx* function with the fragment files and the combined peak set as input. After combining 2-week and adult, the counts were binarized and nuclei having between 4,000 and 25,000 peaks were kept. Furthermore, cells with nucleome signal above 2 and TSS-enrichment score below 0.5 were removed. Peaks were removed if they were present in fewer than 2 cells. Highly variable peaks were selected using *epi.pp.highly_variable* with *min_score=0.53.* Initial Leiden clustering was performed at resolution of 0.5. RNA labels were transferred to the ATAC cells and clustering was refined to yield similar clusters between both omics. Some clusters showed sample-wise separation based on gene expression, which could not be observed in the ATAC. Apart from these RNA-specific subclusters, only those clusters present in both RNA and ATAC were further considered. Differential chromatin openness was also performed using *Scanpy’s sc.tl.rank_genes_groups* on all peaks. Significantly differentially open peaks were defined by having an adjusted p-value below 0.05 and an absolute log_2_-fold change greater than 0.5. In addition to the count matrix on peaks, a count matrix was built similarly on known transcription factor binding sites (TFBS), obtained from https://remap2022.univ-amu.fr/. In this matrix, cells were kept if they had between 4,000 and 50,000 TFBS and TFBS were removed if they were present in fewer than 10 cells. Highly variable TFBS were selected using *epi.pp.highly_variable* with *min_score=0.51.* Cell labels from the gene expression and the peak matrix were transferred to the TFBS matrix and Leiden clustering was performed to annotate the same cell types as for the peak matrix. Finally, the counts were summed up per cell for all binding sites of each unique transcription factor. For the visualization of chromatin openness between cell types at specific regions of interest, we followed the blog post of Andrew John Hill (http://andrewjohnhill.com/blog/2019/04/12/streamlining-scatac-seq-visualization-and-analysis/).

### SCENIC+analyses

To combine the information of gene expression and chromatin openness, SCENIC+ was used (*59*). For this analysis, we followed the SCENIC+ tutorial on analyzing PBMC multiome data from 10X Genomics (https://scenicplus.readthedocs.io/en/latest/pbmc_multiome_tutorial.html#Tutorial:-10x-multiome-pbmc). For the cistopic part of SCENIC+, 48 topics were chosen. In SCENIC+, only eRegulons with a positive correlation between transcription factor expression and target region openness were considered. Visualization was performed as described in the tutorial and network graphs were modified in Cytoscape 3.10.1.

### Mouse scRNA-seq analyses

Annotated sequencing data from scRNAseq in Lyu et al. (*27*) was downloaded from GEO: GSE184175; annotated gene expression data from single cell multiome sequencing in Akagbosu et al. (*34*) was downloaded from GEO: GSE174405. Without any further processing, expression of signature genes for RORγt^+^ DC for the different annotated cell types in each dataset was plotted.

Raw gene expression matrices of scRNAseq in Wang et al. (*38*) were downloaded from GEO: GSE176282 and analyzed using the Seurat package in R. Doublets were identified using the scDblFinder package and subsequently removed. Cells expressing less than 750 unique genes, or with a mitochondrial gene content higher than 5% were also removed. The SCTransform function was used to scale and normalize the data. After dimensional reduction, unsupervised clustering resulted in 23 clusters. Cell types were identified using the genes provided in Fig. S2 of the original publication (*38*) and equivalent populations could be identified. Expression of signature genes for RORC+ DC-like cells was plotted for the different cell types.

Raw gene expression matrices of the CITE-seq dataset in Kedmi et al. (*33*) were downloaded from GEO: GSE200148 and analyzed in R using the Seurat package. Cells from both tdTomato-ON^CD11c^ and tdTomato-ON^RORγt^ x *Zbtb46*-EGFP mice were combined into one workspace. Doublets were removed using the scDblFinder package. Cells with more than 750 unique genes or with a mitochondrial gene content lower than 5% were included for further processing. The SCTransform function was used to scale and normalize the data. After dimensional reduction, unsupervised clustering resulted in 23 clusters which were filtered to only retain cell types included in original publication (*33*). The filtered dataset was re-scaled and dimensional reduction and unsupervised clustering then resulted in 18 clusters, which were identified by the expression of marker genes used for annotation in the original publication (Fig S9e). Expression of signature genes for RORγt^+^ DC was plotted for the different cell types.

### Human scRNA-seq analyses

Raw sequencing data GEO: GSE137710 from Brown et al., Cell, 2019 (*16*) was downloaded. Cells with more than 6000 total RNA fraction and a mitochondrial gene content of less than 7% were isolated. Normalization, scaling, and dimensional reduction of filtered raw counts was performed using the Seurat v4 package. Unsupervised clustering was performed with a resolution of 0.9 resulting in 11 clusters which were identified by the expression of marker genes. Cells annotated as CD1c^+^ myeloid dendritic cells, CD141^+^ myeloid dendritic cells, innate lymphoid cell and plasmacytoid dendritic cells from the spleen scRNA-seq data from the Tabula Sapiens Consortium (*62*) were reclustered. Normalization, scaling, and dimensional reduction was performed using Seurat v4. Unsupervised clustering at resolution 1.4 resulted in 7 clusters, which were classified by the expression of marker genes.

The published normalized and log-transformed scRNA-seq data from the Gut Cell Atlas (*65*) was analyzed using *Scanpy*. Cells annotated as derived from healthy adult donors and annotated as lymphoid DC, cDC1, cDC2 and pDC were selected for further analysis. After dimensionality reduction unsupervised clustering was performed at a resolution of 1.5. The resulting cluster 0, which contained XCR1^+^ cDC1 and ZFP36L2^+^ cDC2 was subclustered at a resolution of 0.4 to better reflect the DC subset heterogeneity. The resulting 13 clusters were identified by expression of marker genes. Cells derived from paediatric healthy donors or donors with Crohn’s diagnosis annotated asl DC, cDC1, cDC2 and pDC were selected for further analyses. Post dimensionality reduction, unsupervised clustering was performed at a resolution of 1.6. The resulting 13 clusters were identified by expression of marker genes.

The Tabula Sapiens Lymph Node dataset {Ref} was analysed using *Scanpy*. After normalization and log-transformation of the data, cells annotated as CD1c^+^ myeloid DC, CD141^+^ myeloid DC, conventional DC, hematopoietic stem cell, innate lymphoid cell and plasmacytoid DCs were selected for further analysis. After performing dimensionality reduction, unsupervised clustering at a resolution of 1.0 resulted in 15 clusters which were identified by expression of marker genes.

After pre-processing in *Scanpy* the above datasets were converted to Seurat objects and further processed using Seurat v4 package. Genes expressed in more than 20% of cells and with an adjusted p-value <0.05 and log2_FC > 0.4 were considered for differential gene expression analyses. To score cells for enrichment of specific gene signatures, the AUCell package was used. Signatures from the mouse single cell multiome analyses were converted to human orthologs using the Orthogene package prior to scoring. Integration of the two human spleen datasets was performed according to the standard Seurat Integration workflow. Both datasets were normalized by SCTransform before finding integration features and integration anchors. Subsequently, dimensional reduction was performed to display cells from both datasets in the same low-dimensional space.

### Neonatal murine small intestinal scRNA-seq analyses

Sequencing data were processed using 10X Genomics Cell Ranger v6.0.0 pipeline using default parameters. The resulting count matrix was loaded into R (v4.3.2) and downstream analyses were performed using Seurat v4 and v5. We removed genes detected in less than 3 cells from further analysis and only included cells with less than 5% mitochondrial gene content, or between 1500 and 6000 different genes detected per cell. Hash Tag Oligo (HTO) information was integrated and HTO demultiplexing was performed per Seurat workflow, excluding cells marked as doublets. A second round of doublet exclusion was performed using the DoubletFinder package (https://github.com/chris-mcginnis-ucsf/DoubletFinder). The resulting Seurat object contained 1391 cells consisting of 662 cells belonging to P1 timepoint and 729 cells belonging to the P9 timepoint. Cell cycle state was scored based on published cell cycle gene lists (https://satijalab.org/seurat/articles/cell_cycle_vignette). SCTransform package was used for scaling, normalization, and cell cycle regression. The first 60 Principal Components were used for dimensional reduction using UMAP. Unsupervised clustering was performed with a resolution of 1.5. *FindAllMarkers* function of Seurat was used to calculate differentially expressed genes between the clusters.

### Integration of datasets

The indicated published datasets were downloaded from GEO: GSE184175, GSE174405, GSE176282, and GSE200148 and the original cell type annotation used except for GSE200148 (*33*) and GSE176282 (*38*), which were annotated analogous to the original publications as described above. For easier integration and higher comparability between datasets we grouped cell types into broad categories, e.g. subgroups of T-cells only described in one dataset (*27*) were combined into one T-cell cluster, B cell subgroups into a B cell cluster and so forth. Furthermore, cell types that were present in only one of the datasets were removed to facilitate the integration task. The raw datasets were combined into an AnnData object. On the combined data, cells were included if they had between 1000 to 50,000 counts and if they had between 500 genes and 6000 genes. Any cells outside these ranges were excluded. Genes were removed if they were detected in under 5 cells. Scanpy’s *sc.pp.highly_variable_genes* was used with the datasets as batch variable to select the top 2000 variable genes for integration. Finally, scVI was used for unsupervised integration, following the tutorial https://docs.scvi-tools.org/en/stable/tutorials/notebooks/scrna/harmonization.html. For visualization, UMAPs were calculated on the scVI embedding. Additionally, Leiden clustering was performed on the scVI embedded cells at resolution 0.2, resulting in 10 clusters closely corresponding to the originally annotated broad cell types across datasets (exceptions: ILC2 clustered with T-cells, LTi-ILC3 and ILC3 were together in one cluster, proliferating DC clustered with pDC, and Mac clustered with cDC2). As shown in Fig. 6d, the “RORγt^+^ DC” cluster contained *Rorc*^+^ or *Aire^+^* APC populations annotated distinctly in various publication as well as the RORγt^+^ DCs and RORγt^+^ eTACs reported in this publication. To further validate how well the Leiden clusters represented the reported cell types, the percentage of cells from a given cell type in each Leiden cluster was calculated and visualized in a heatmap. Additionally, the ARI score between the broad cell type annotation based on the original publications and the Leiden clusters on the integrated datasets was calculated. This resulted in a score of 0.64.

### Statistical analyses

Statistical significance was calculated in Prism 10 software (GraphPad). For pairwise comparisons two-tailed t-test with Welch’s correction was used. For multiple comparisons, one-way analysis of variance with Tukey’s test was performed. Wilcoxon non-parametric ranked sum test was used to calculate differences in AUCscores in R. A p-value <0.05 was considered significant.

## Acknowledgements

We thank Anne Krug and members of the Schraml lab for helpful discussions and critical reading of the manuscript. We also thank members of the Krug and Dudziak laboratories and Malte Benjamin Braun from the Renkawitz laboratory for technical help. We thank Vasileios Bekiaris for providing *RORγt^GFP^* mice for migration experiments. We acknowledge the Core Facilities for Flow Cytometry, for Bioimaging and for Animal Models at the Biomedical Center, LMU Munich for providing equipment and expertise. High-throughput sequencing was performed by the laboratory for Functional Genome Analysis (LAFUGA) of LMU Munich.

## Author contributions

H.N. and B.U.S. designed experiments. H.N. performed experiments with help from N.E.P., C.S., I.U., K.R.R., and V.K. M.L.R., R.S. and K.R.R. performed bioinformatic analyses with help from E.L.S. and A.U.A. N.E.P. first observed that RORγt^+^ DCs exist in adult spleen and non-lymphoid organs. D.D. provided targeting antibodies. L.K. provided *Aire-hCD2* mice, C.R. provided *Rag2*^-/-^*γc*^-/-^ and *RORγt^Cre^Rosa^YFP^RORγt^GFP^* mice, and both helped in planning the corresponding experiments. M.C.T. and M.L.R. provided critical input for single cell ATAC and scRNA-seq experiments and analyses. K.L., M.C.T., C.R., D.D., N.T., M.W.H., M.C. and L.K. provided reagents and critical intellectual input. B.S. designed and supervised the study. B.S. and H.N. wrote the manuscript with input from all authors.

## Data and materials availability

The authors declare that the data supporting the findings of this study are available within the paper and its supplementary files. Raw data are available from the authors upon reasonable request. Datasets related to single-cell RNA sequencing and multiome experiments that were generated and analyzed for the current study have been deposited at BioStudies database under accession number S-BSST1322.

---

**Supplementary Figure 1.**
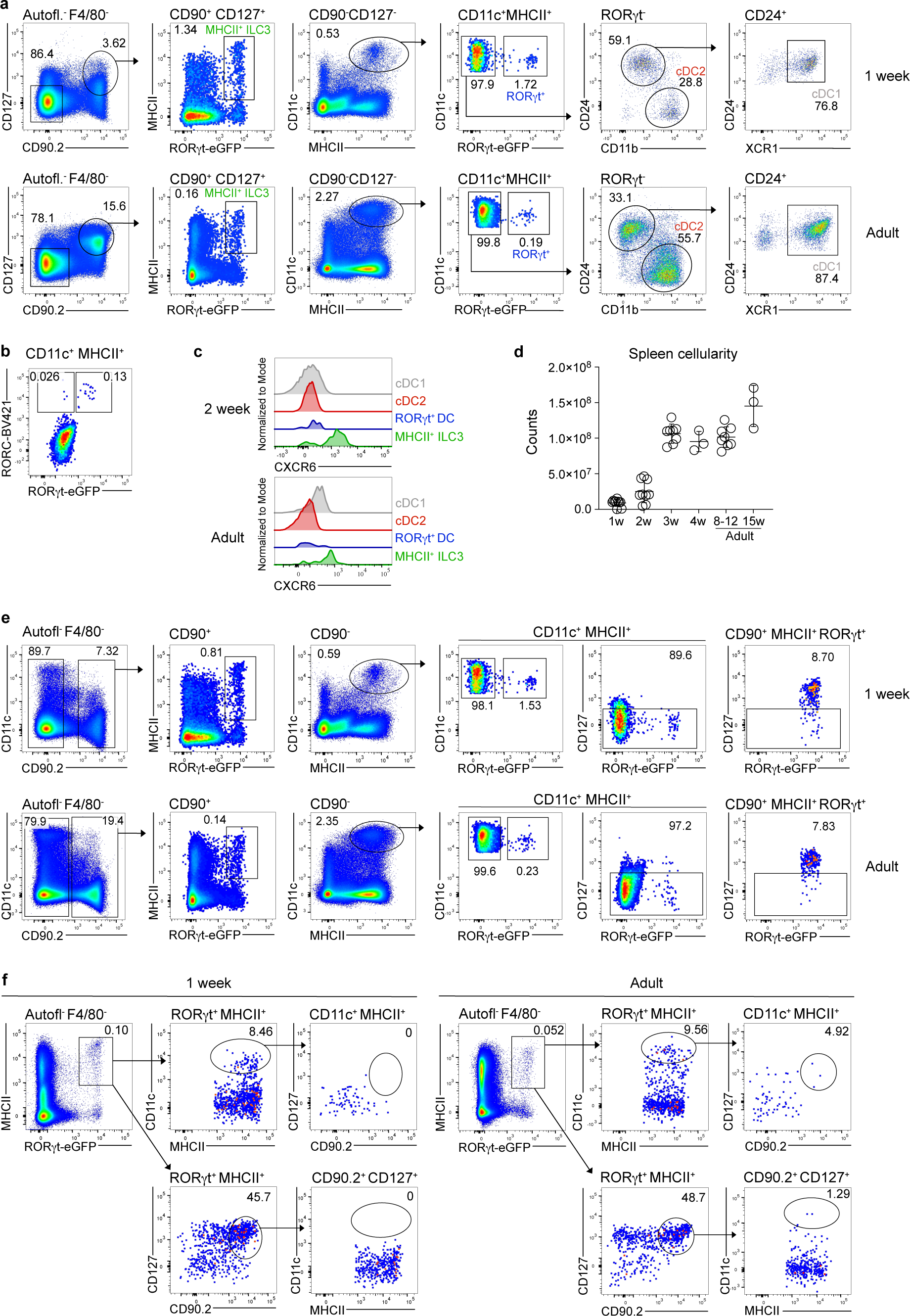
CD11c^+^MHCII^+^RORγt^+^ cells are phenotypically more similar to cDCs than MHCII^+^ILC3 and conserved across age. **a-f.** Spleens from *RORγt^GFP^Clec9a^Cre^Rosa^Tom^*or *Clec9a^Cre^Rosa^Tom^* mice at the indicated ages were analyzed by flow cytometry **a.** Representative gating strategy for cDC1, cDC2, MHCII^+^ILC3 and RORγt^+^ DC. Live autofluorescence negative cells were gated. Within CD127^+^CD90^+^ cells MHCII^+^GFP^+^ cells were gated as MHCII^+^ ILC3. In the CD127^-^CD90^-^ fraction CD11c^+^MHCII^+^ cells were gated and RORγt^+^ DC identified as GFP^+^ or RORC positive cells **(b)**. CD11c^+^MHCII^+^GFP^-^ cells were further divided into CD11b^+^ cDC2 and CD24^+^ cells. CD24^+^XCR1^+^ cells were identified as cDC1. **b.** CD11c^+^MHCII^+^ cells from *RORγt^GFP^* mice were stained using an anti-RORC antibody to demonstrate that anti-RORC staining and RORγt driven GFP are largely congruent. **c.** Representative histogram overlays showing expression of CXCR6 in cDC1 (grey), cDC2 (red), CD11c^+^MHCII^+^ RORγt^+^ cells (blue) and MHCII^+^ ILC3 (green) from 2-week-old and adult mice. **d.** Spleen leukocyte cellularity at the indicated ages. Each dot represents one mouse **e.** CD11c^+^MHCII^+^RORγt^+^ cells do not express CD127. CD90^-^ CD11c^+^MHCII^+^ cells were gated without prior exclusion of CD127^+^ cells and then analyzed for GFP and CD127 expression. CD90^+^MHCII^+^GFP^+^ ILC3 stained positive for CD127. **f.** RORγt^+^ MHCII^+^ cells in 1-week-old and adult spleen were gated and further analyzed for expression of CD11c, CD90 and CD127. Data are representative of at least two independent experiments with at least 3 mice per group.

**Supplementary Figure 2.**
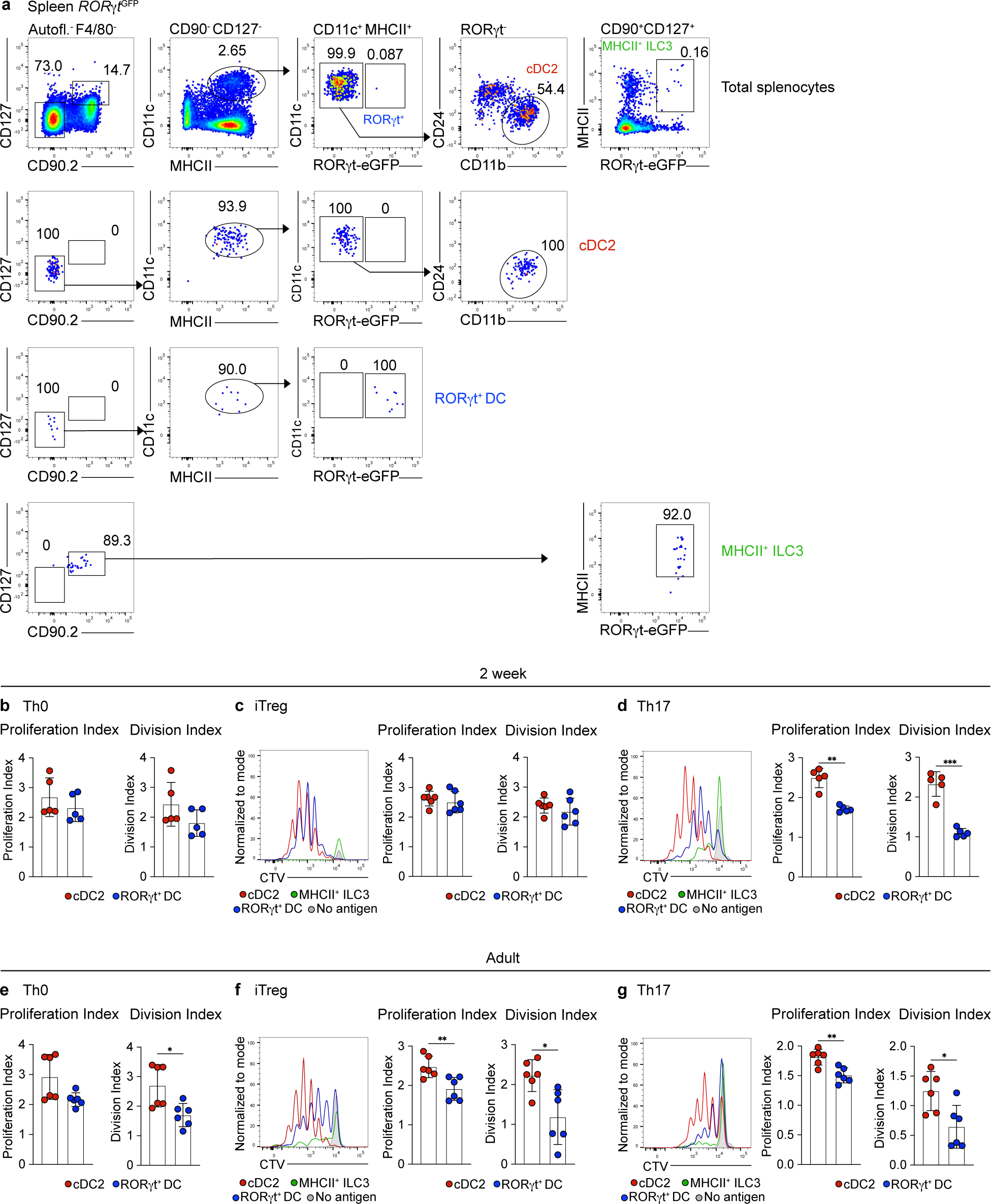
Sort purity of APC populations and proliferation of OTII cells. **a-g.** cDC2, MHCII^+^ ILC3 and GFP^+^ RORγt^+^ DC were sorted from spleens of 2-week-old or adult *RORγt^GFP^* mice and pulsed with OVA_323-339_. 250 APCs were then co-cultured with 2500 CTV-labelled naïve OTII cells under the indicated conditions for 3.5 days. **a.** Gating strategy (top row) and sort purity analyses of cDC2, MHCII^+^ ILC3 and RORγt^+^ DC from adult mice. **b-g.** Division and proliferation indexes of OTII cells after co-culture with the specified populations from 2-week-old **(b-d)** or adult **(e-g)** mice under indicated conditions. Dots represent biological replicates from at least 2 independent experiments. *p (0.0332), **p (0.0021) ***p (0.0002), ****p < 0.0001. Statistical analysis was performed using two-tailed Welch’s t-test. Only statistically significant comparisons are indicated.

**Supplementary Figure 3.**
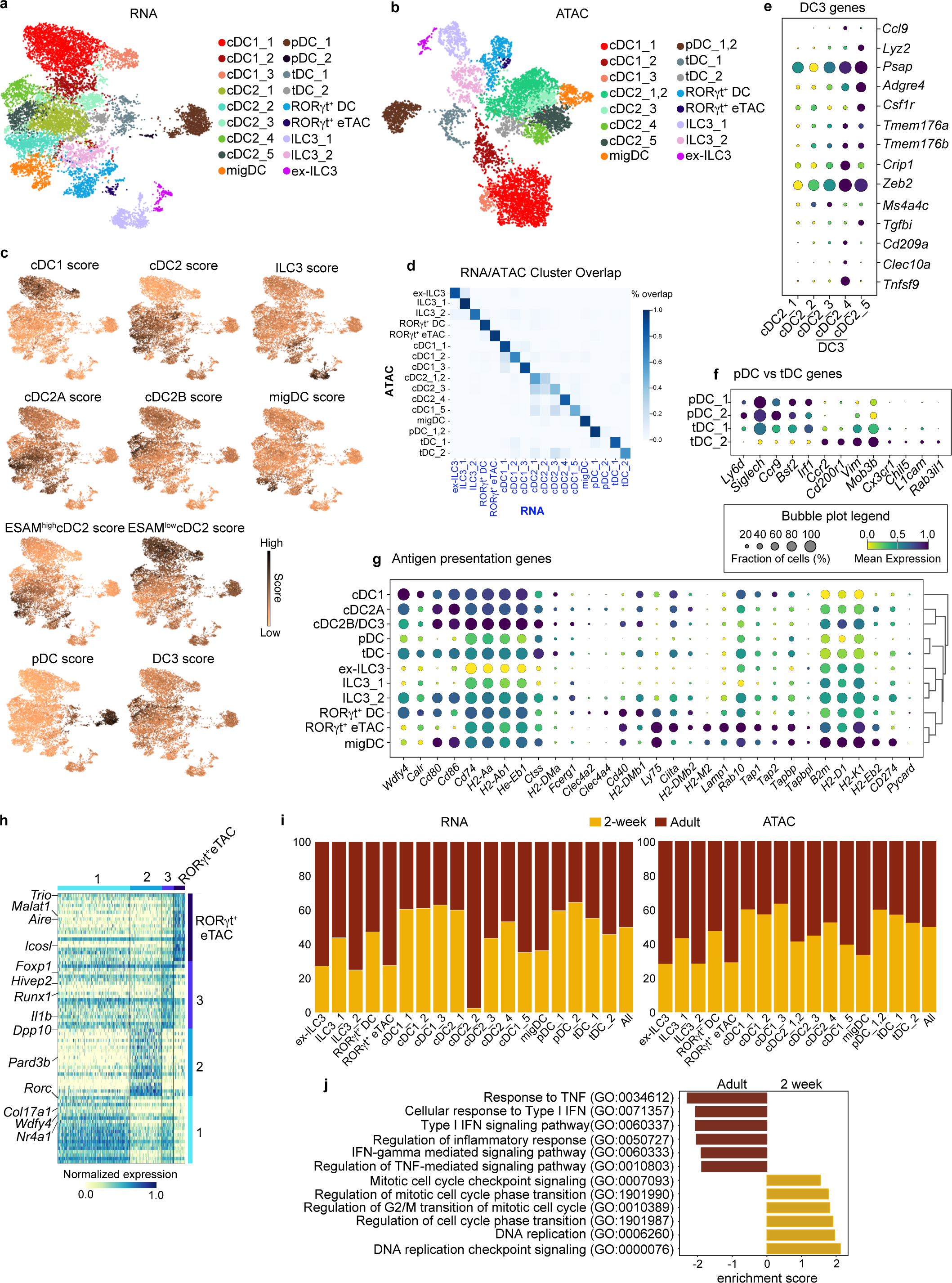
Cluster annotation and downstream analyses of single cell multiomics profiling. RORγt^+^ DC, CD11c^+^MHCII^+^ cDC and MHCII^+^ ILC3 from spleens of 2-week-old (n=2) or adult (n=3) *RORγt^GFP^Clec9a^Cre^Rosa^Tom^* mice were sorted, pooled at a 1:10:1 ratio and subjected to 10X multiome analyses. **a.** Annotated UMAP representing RNA-based analysis of 11,980 nuclei. **b.** Annotated UMAP representing open chromatin based analysis of 9899 nuclei. **c.** RNA-based UMAPs depicting expression scores for the indicated gene signatures provided in Table 1. **d.** Correspondence of clusters between scATAC and scRNA based analyses. **e-g**. Bubble plots showing mean expression of select DC3 genes **(e)**, pDC and tDC genes **(f)**, and for genes involved in antigen presentation **(g)**. **h.** Heatmap showing top 20 differentially expressed genes between RORγt^+^ DC clusters and RORγt^+^ eTACs with selected genes highlighted. For full list of genes see Table 4. **i.** Contribution of cells from the 2-week (yellow) or adult (red) timepoints to RNA and ATAC based clusters. **j.** Normalized enrichment scores of selected gene sets enriched in all cDC2 clusters from 2-week-old and adult mice.

**Supplementary Figure 4:**
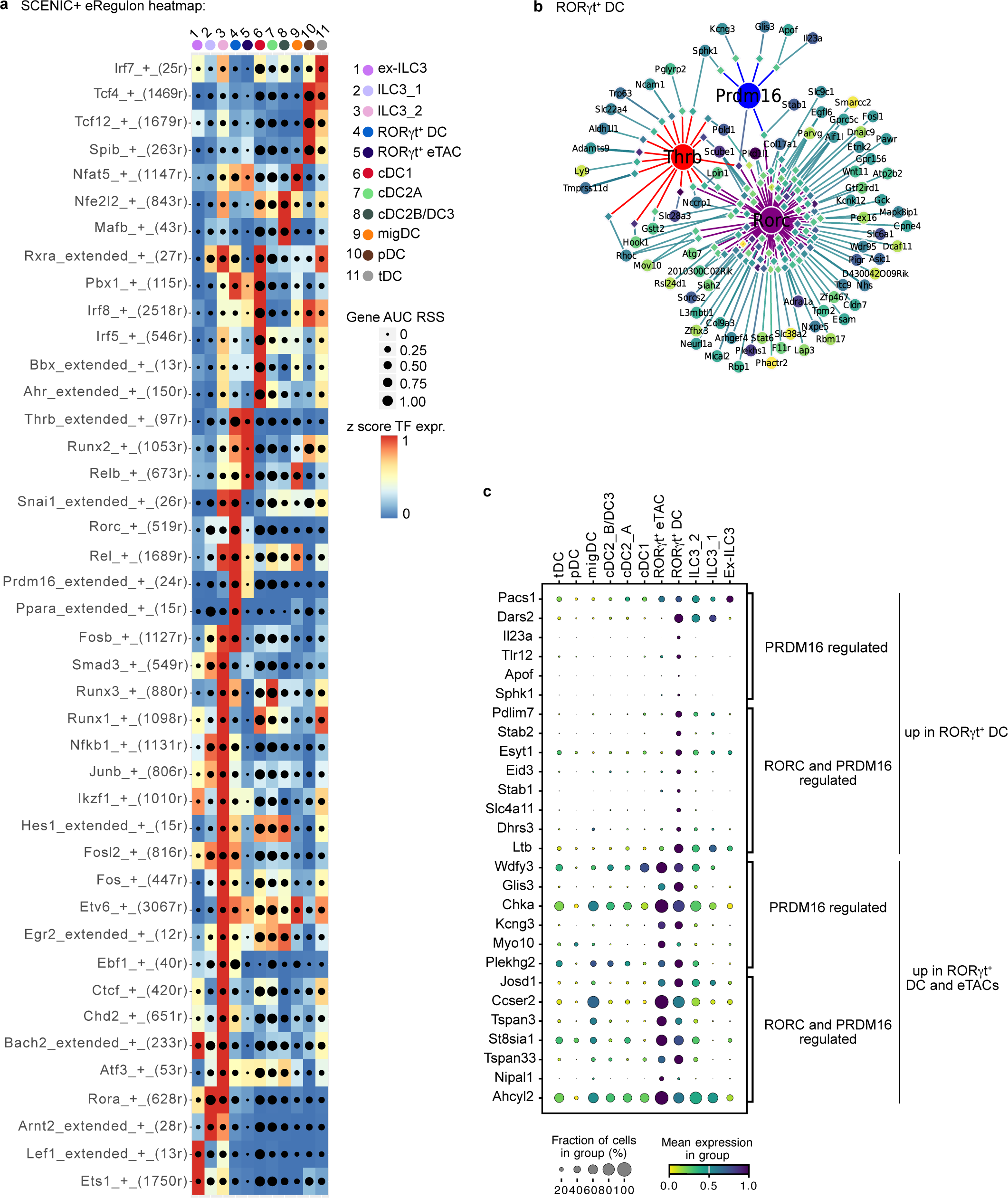
Integrated multiomic analyses using SCENIC+ reveals gene regulatory networks distinguishing RORγt^+^ DC. **a.** Heat map/dot-plot showing eRegulons computed with SCENIC+. Transcription factor expression is given on a color scale and cell-type specificity of the genes regulated by the transcription factors (eRegulon) on a size scale. **b.** Visualization of the gene regulatory network formed by *Prdm16*, *Thrb* and *Rorc* in RORγt^+^ DC. **c.** Bubble plot of genes regulated by PRDM16 or PRDM16 and RORγt in RORγt^+^ DCs or RORγt^+^ eTACs.

**Supplementary Figure 5.**
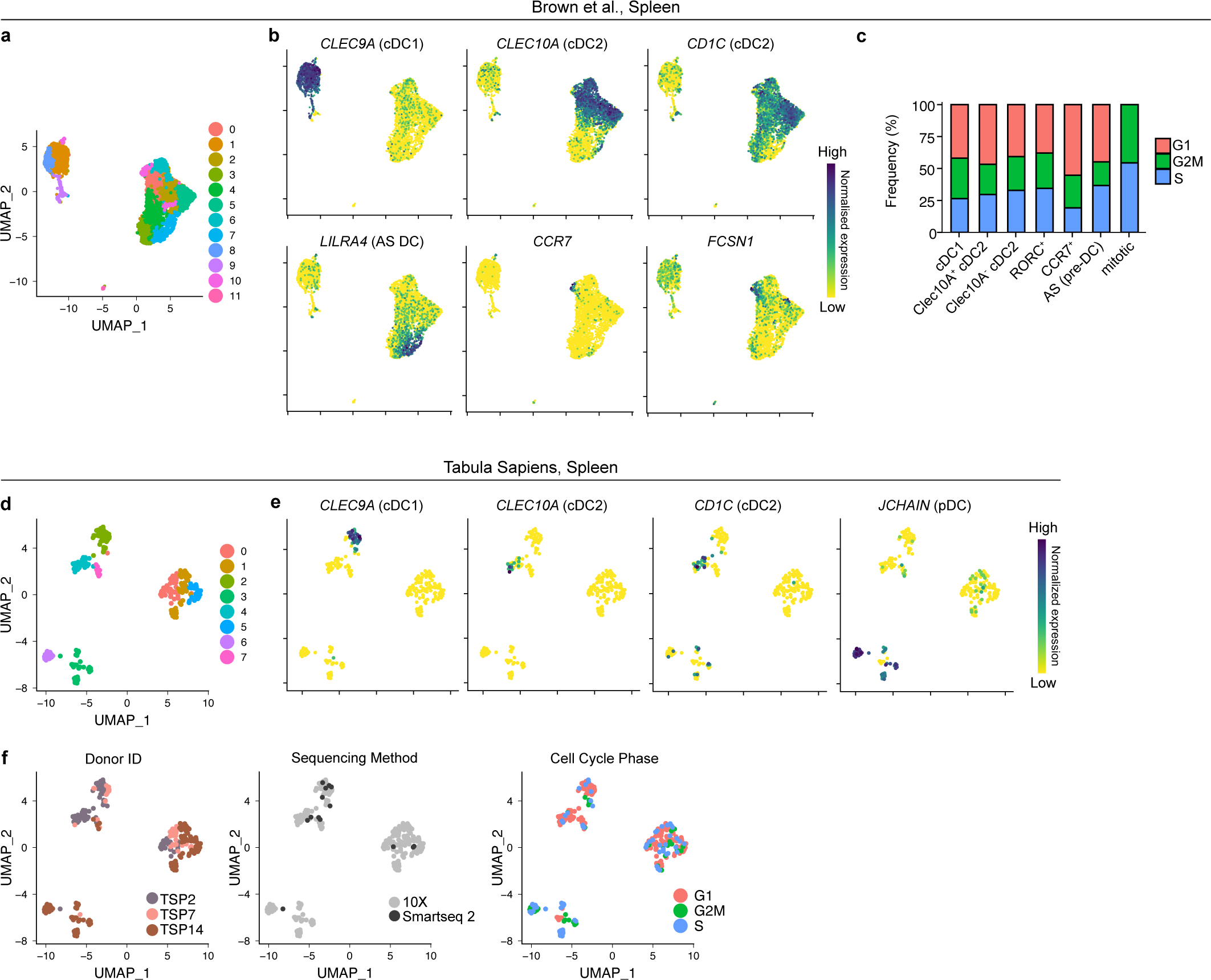
RORγt^+^ DC exist in human spleen. **a-c**. Human splenic DCs profiled by scRNASeq in Brown et al. were analyzed using Seurat algorithm. **a.** UMAP display of 4717 cells colored by cluster and **b.** Expression of genes used for cluster annotation. **c**. Proportion of cells per annotated cluster in indicated cell cycle phases. **d-f.** 262 cells annotated as DCs, pDCs and ILCs in the scRNAseq spleen reference dataset from the Tabula Sapiens Consortium were reclustered. UMAP displaying Leiden clustering **(d)** and expression of the indicated genes used for cluster annotation **(e)**. **f.** UMAP annotated by donor ID (left), sequencing method (middle) and cell cycle phase (right).

**Supplementary Figure 6.**
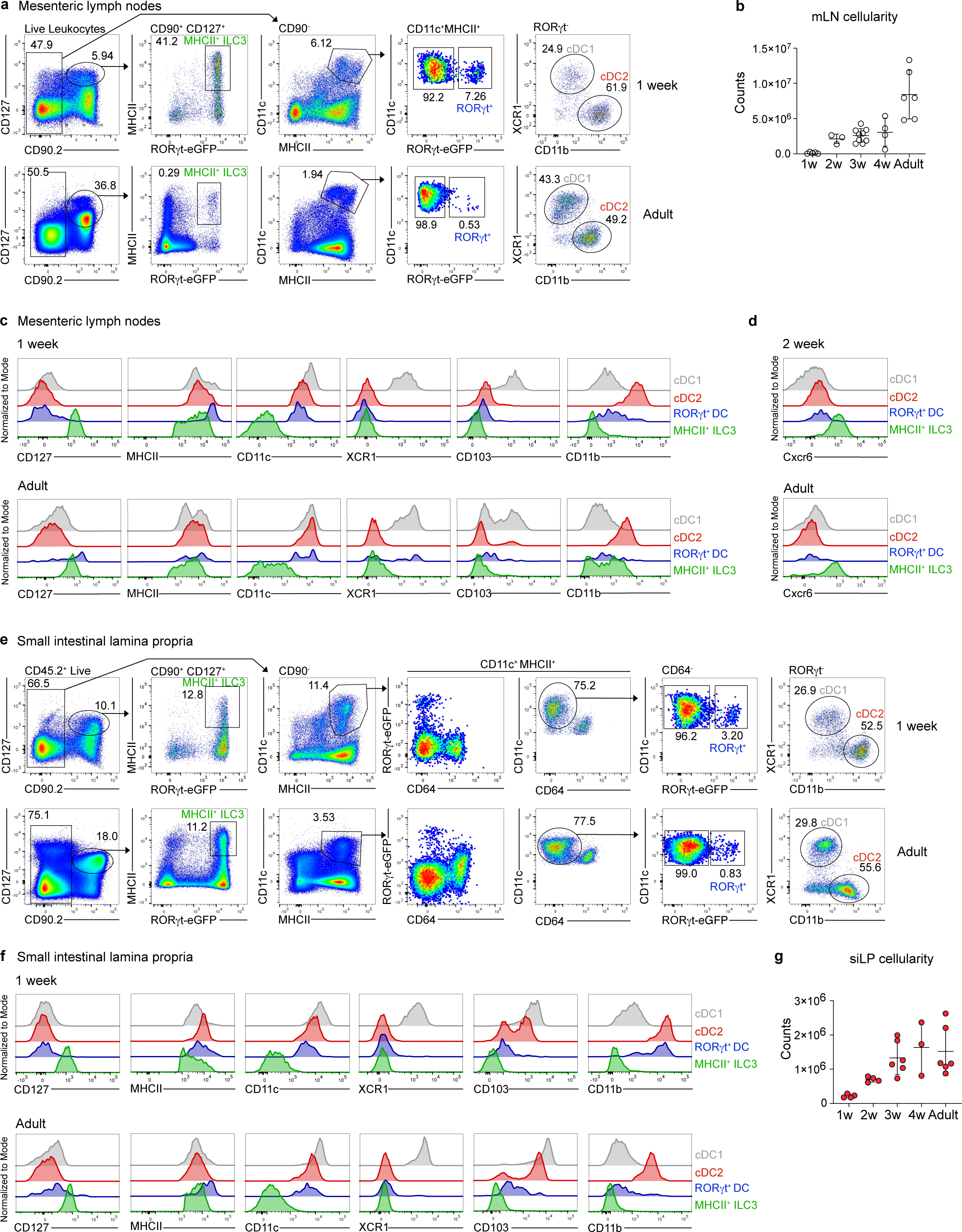
Characterization of RORγt^+^ DC in mesenteric lymph node and small intestinal lamina propria of neonatal and adult mice. Mesenteric lymph node (mLN) **(a-d)** and small intestinal lamina propria (siLP) **(e-g)** from *RORγt^GFP^* mice at the indicated ages were analyzed by flow cytometry. **a.** Representative gating strategy for the identification of MHCII^+^ ILC3, RORγt^+^ DC, cDC1 and cDC2 in mLN of 1-week-old and adult mice. **b.** mLN cellularity across age (1 week n=5; 2 week n=3; 3 week n=8, 4 week n=4; Adult n=6). **c, d.** Histogram overlays of the indicated surface markers on cDC1 (grey), cDC2 (red), RORγt^+^ DC (blue) and MHCII^+^ ILC3 (green). **e.** Representative gating strategy for identification of MHCII^+^ ILC3, RORγt^+^ DC, cDC1 and cDC2 in siLP of 1-week-old and adult mice. **f**. Histogram overlays of the indicated markers on cDC1 (grey), cDC2 (red), RORγt^+^ DC (blue) and MHCII^+^ ILC3 (green). **g.** Total leukocyte count per siLP at the indicated ages (1-week-old n=4-8; 2-week-old n=4; 3-week-old n=6, 4-week-old n=3; Adult n=6). mLN from 1-week-old mice were pooled by 2 or 3, otherwise each dot represents one mouse. Error bars represent mean ± SD.

**Supplementary Figure 7.**
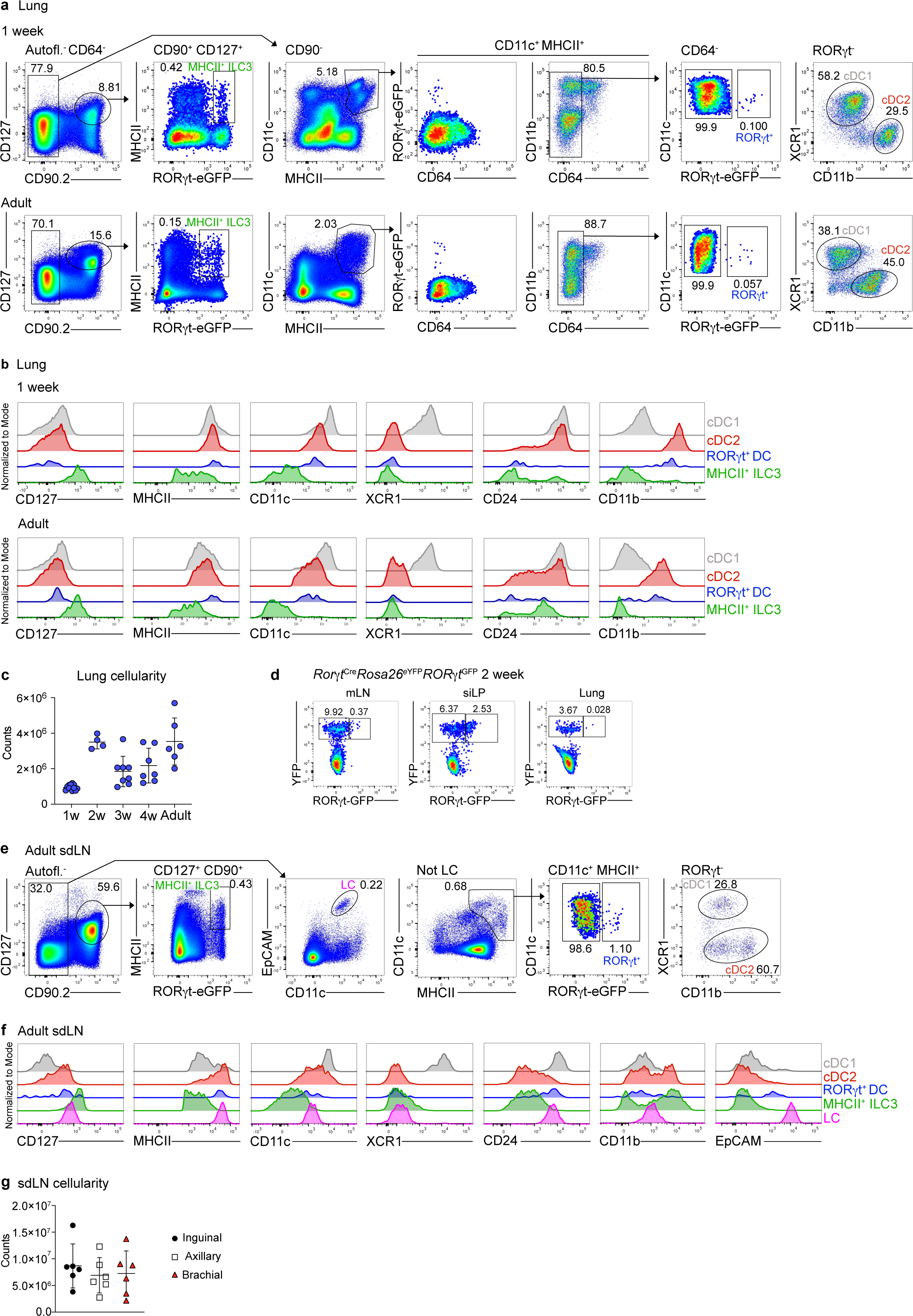
Identification and phenotypic characterization of RORγt^+^ DC in murine lung and skin draining lymph nodes. **a.** Representative gating strategy for the identification of MHCII^+^ ILC3, RORγt^+^ DC, cDC1 and cDC2 in murine lung at indicated ages. **b.** Histogram overlays of the indicated surface markers on cDC1 (grey), cDC2 (red), RORγt^+^ DC (blue) and MHCII^+^ ILC3 (green). **c.** Lung cellularity across age (1-week-old n=11; 2-week-old n=4; 3-week-old n=7, 4-week-old n=6; Adult n=6) from mice of indicated ages. Data are pooled from at least 2 independent experiments. Each dot represents one mouse. **d.** mLN, siLP and lung from 2-week-old *RORγt*^Cre^*Rosa*^YFP^*RORγt*^GFP/wt^ mice were analyzed by flow cytometry. CD11c^+^MHCII^+^ cells were gated as indicated above and analyzed for GFP (active RORγt) versus YFP (RORγt expression history). **e-g.** Axial, brachial, and inguinal sdLN were isolated from adult *RORγt^GFP^* mice. **e.** Representative gating strategy for identification of MHCII^+^ ILC3, Langerhans cells (LC), RORγt^+^ DC, cDC1 and cDC2 in adult murine adult skin-draining lymph nodes. **f.** Histogram overlays of the indicated surface markers on cDC1 (grey), cDC2 (red), RORγt^+^ DC (blue), MHCII^+^ ILC3 (green) and LC (pink). **g.** Axial, brachial, and inguinal sdLN cellularity for each site. Data pooled from 2 independent experiments performed with 3 mice each, each dot represents a biological replicate. Error bars represent mean ± SD.

**Supplementary Figure 8.**
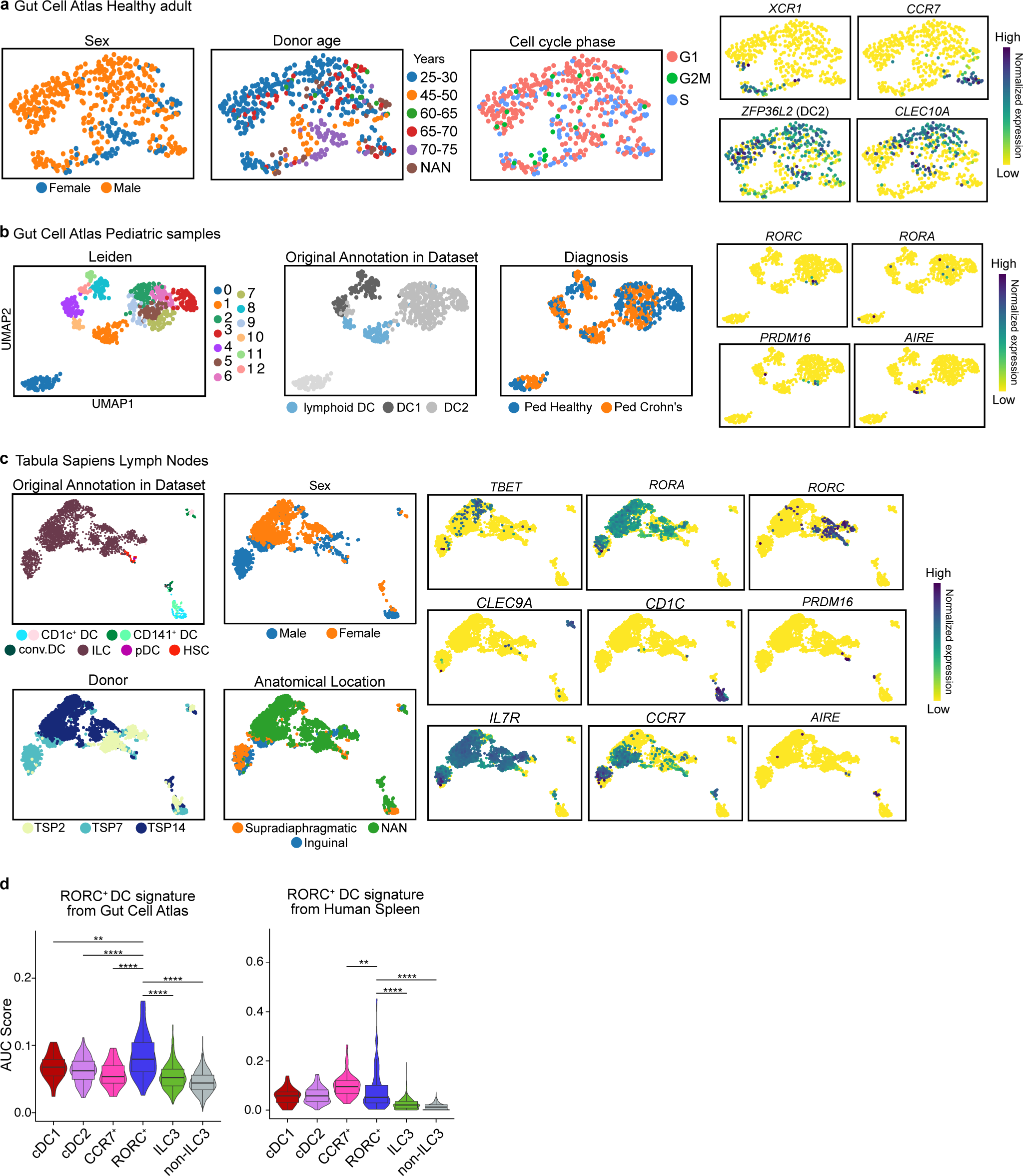
RORγt^+^ DC in scRNAseq datasets of human tissues. **a, b**. Human gut cell atlas single cell transcriptome analyses. **a.** 480 cells annotated as lymphoid DCs, DC1 and DC2 DCs from healthy adult donors from the human gut cell atlas were reclustered using *Scanpy*. UMAP display depicting sex, donor age, cell cycle phase and the indicated genes used for cluster identification in Fig. 5e. **b.** 644 cells annotated as lymphoid DC, DC1 and DC2 from pediatric donors from the human gut cell atlas were reclustered using *Scanpy* into 13 clusters as indicated on the left UMAP. Middle UMAPs depict the original cluster annotation and stratification of cells per donor diagnosis (healthy vs. pediatric Crohn’s disease). UMAPs on the right show expression of the *PRDM16, RORC, AIRE*, and *RORA* to visualize presence of *PRDM16^+^RORC^+^*cells resembling RORγt^+^ DC, distinct from AIRE-expressing cells. **c.** 2668 cells from the Tabula Sapiens Lymph node reference data set originally annotated as CD1c^+^DC, CD141+DC, cDC (conv.DC), ILC, pDC, and HSC were reclustered using *Scanpy*. The resulting UMAP is colored by original cell annotation, sex, donor, anatomical location, and the indicated marker genes used for cluster identification. **d.** Enrichment score for expression of genes distinguishing RORγt^+^ DC in the healthy adult gut (left) and healthy adult spleen (right, Fig. 4) across annotated clusters in Tabula Sapiens Lymph Nodes dataset. *p (<0.05), **p (<0.01) ***p (<0.001), ****p < 0.0001. Statistical analysis was performed using Wilcoxon non-parametric ranked sum test to calculate differences in AUCscores between indicated clusters.

**Supplementary Figure 9.**
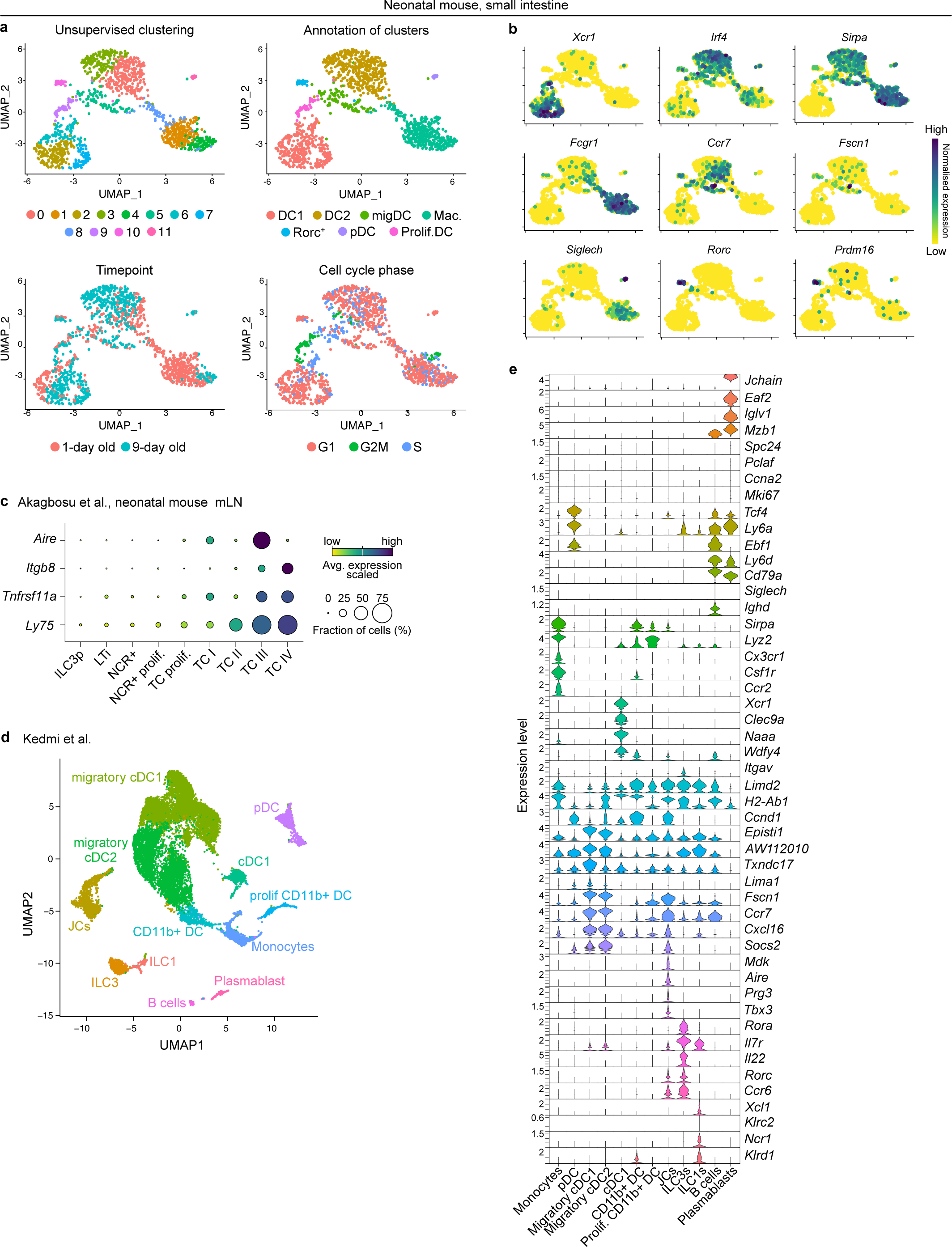
RORγt^+^ DC in scRNAseq datasets of murine tissues. **a, b.** scRNAseq of neonatal siLP. Live CD45^+^CD19^-^CD90^-^CD11c^+^MHCII^+^ cells from siLP of one-day old and nine-day old *Clec9a^Cre^Rosa^Tom^* mice were sorted, pooled, and subjected to 10X scRNAseq analyses. **a.** UMAP display of 2053 cells colored by cluster, cell type annotation, timepoint and cell cycle phase. **b.** UMAP showing expression of the indicated genes used for cluster annotation. **c.** Expression of the indicated genes in clusters from Akagbosu et al. **d.** UMAP display of 16,302 cells in dataset from Kedmi et al. colored by clusters. **e.** Stacked violin plots for selected genes from Kedmi et al. Extended Data Fig 2.

**Supplementary Figure 10.**
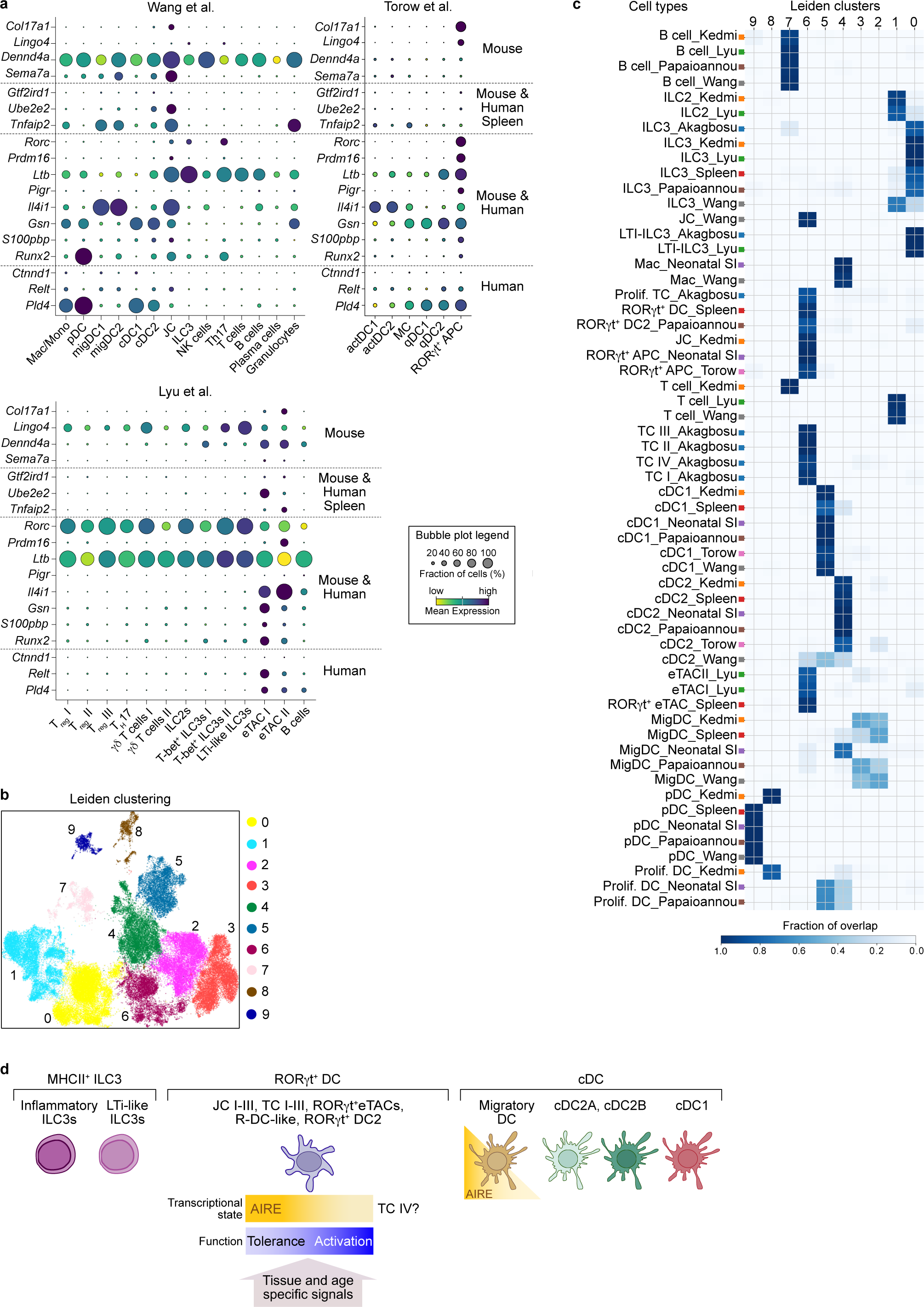
RORγt^+^ DC include previously described RORγt^+^ APC subtypes. **a.** Bubble plots showing expression of the indicated genes in the specified murine scRNAseq datasets. Genes are ordered according to the species and organs they were deduced from. **b.** UMAP display of integrated murine datasets from Fig. 6b colored by Leiden clusters. **c.** Heatmap depicting fraction of overlap of cell types from the indicated datasets with each Leiden cluster. **d.** RORγt^+^ DC are a distinct immune lineage that entails Janus cells, Thetis cells, RORγt^+^ eTACs, R-DC-like cells and RORγt^+^ DC2. RORγt^+^ DC are transcriptionally heterogeneous and exhibit a spectrum of *Aire* expression, captured in our analysis and previous work describing substates of Janus cells (JCI-III) and Thetis cells (TCI-IV). We speculate that this heterogeneity reflects a functional spectrum, ranging from inducing T cell activation to tolerance. These transcriptional states and functions of RORγt^+^ DCs maybe shaped by tissue and age-specific signals.

